# Biochemical properties of chromatin domains define genome compartmentalization

**DOI:** 10.1101/2024.03.05.583467

**Authors:** Federica Lucini, Cristiano Petrini, Elisa Salviato, Koustav Pal, Valentina Rosti, Francesca Gorini, Philina Santarelli, Roberto Quadri, Giovanni Lembo, Giulia Graziano, Emanuele Di Patrizio Soldateschi, Ilario Tagliaferri, Eva Pinatel, Endre Sebestyén, Luca Rotta, Francesco Gentile, Valentina Vaira, Chiara Lanzuolo, Francesco Ferrari

## Abstract

Chromatin three-dimensional (3D) organization inside the cell nucleus determines the separation of euchromatin and heterochromatin domains. Their segregation results in the definition of active and inactive chromatin compartments, whereby the local concentration of associated proteins, RNA and DNA results in the formation of distinct subnuclear structures. Thus, chromatin domains spatially confined in a specific 3D nuclear compartment are expected to share similar epigenetic features and biochemical properties, in terms of accessibility and solubility.

Based on this rationale, we developed the 4f-SAMMY-seq to map euchromatin and heterochromatin based on their accessibility and solubility, starting from as little as 10,000 cells. Adopting a tailored bioinformatic data analysis approach we reconstruct also their 3D segregation in active and inactive chromatin compartments and sub-compartments, thus recapitulating the characteristic properties of distinct chromatin states.

A key novelty is the capability to map both the linear segmentation of open and closed chromatin domains, as well as their 3D compartmentalization in one single experiment.

## INTRODUCTION

In the interphase cell nucleus, chromatin is organized in a three-dimensional (3D) architecture reflecting the epigenetic and transcriptional regulation of the genome (1). Alterations of chromatin 3D architecture have been identified in multiple human diseases (2–4). On a large-scale, euchromatic (active) and heterochromatic (inactive) domains tend to co-localize with domains of the same type, thus constituting spatially separated compartments that can be mapped by high-throughput genome-wide chromosome conformation capture (Hi-C) (5). The active and inactive compartments are conventionally named “A” and “B” compartments, enriched in transcribed or repressed genomic regions, respectively. The Nuclear Lamina (NL) contributes to this compartment separation by binding specific heterochromatic genomic regions called Lamina Associated Domains (LADs) (6), belonging to the “B” compartment. Over the years, improvements in the resolution of experimental data (7) and in the computational analyses (8) allowed identifying also sub-compartments, whereby a finer grain segmentation of “A” and “B” compartments can be associated to specific combinations of chromatin marks, thus achieving a more precise link between the 3D chromatin organization and epigenetic regulation. Additional methodological improvements in this field focused on achieving a higher resolution in mapping contacts between genomic loci, e.g. with Micro-C (9,10), or a reduced number of starting cells, e.g. with Low-C (11). Although undoubtedly powerful, Hi-C is based on a multistep protocol including chemical modifications and PCR amplification that can contribute technical biases masking or underestimating small chromatin remodelling dynamics. Moreover, by measuring the pairwise contact frequency of genomic loci, it does not give information on whether a specific contact is occurring in a particular subnuclear region. To this concern, multiple pieces of evidence in literature highlighted that the detachment of chromatin from lamina causes a major reorganization of chromatin 3D compartmentalization that is visible by imaging techniques, but does not result in evident changes in Hi-C compartments (12). In a different model with triple knock-out of lamina proteins, also the Topologically Associating Domains (TADs) detected by Hi-C have been shown to be largely unchanged (13). Indeed, as long as the local contact pairs are preserved, the chromosome conformation capture-based techniques may not be able to detect the change of location of a domain across subnuclear regions. Alternative methods to overcome some technical limitations of ligation by proximity were proposed with GAM (14) and SPRITE (15) techniques. Nevertheless, these approaches, that require highly specialized instruments and expertise, are also based on the detection of contact frequencies between genomic loci. We recently presented SAMMY-seq, that is based instead on the biochemical fractionation of chromatin, followed by sequencing of the individual fractions to map the constitutive heterochromatin domains. We already reported that SAMMY-seq is able to detect early changes in heterochromatin solubility when LAD association to the nuclear lamina is compromised due to a mutated form of Lamin A (16). Notably, in the same early passage cellular model, Hi-C could not detect 3D chromatin architecture changes (17). More recently, in a completely different model of mechanical stress induced by cell motility, where the deformation of cell nucleus is expected to trigger a dissociation of heterochromatic LADs from the lamina (18), SAMMY-seq could detect reorganization dynamics involving heterochromatin domains (19). Despite the sensitivity in detecting heterochromatin dynamics, the original SAMMY-seq method had limited resolution in mapping sharp localized peaks of chromatin accessibility, for example those associated with active gene promoters.

Here we present a novel experimental protocol (4f-SAMMY-seq) and dedicated data analyses algorithms to map the position of both open and closed chromatin regions along the genome, in addition to their 3D spatial segregation in distinct chromatin compartments. These innovations with respect to earlier SAMMY-seq are achieved with two novel steps in the experimental protocol, including a lighter partial digestion of accessible chromatin and a size separation step to map more precisely the location of sharp peaks of accessibility. Moreover, a novel analysis procedure allows reconstructing chromatin segmentation in spatially confined compartments and sub-compartments with a level of details previously achieved only with Hi-C. To this concern, 4f-SAMMY-seq provides crucial practical advantages over state-of-the-art techniques for mapping compartments in terms of costs, versatility and scalability. Indeed, the novel 4f-SAMMY-seq works on as little as 10,000 (10k) cells, it requires only a few hours of bench work and a limited sequencing depth, as low as 25 million reads per chromatin fractions. These advantages open unprecedented possibilities for characterizing chromatin 3D compartmentalization and scaling up its analysis in a variety of experimental settings.

## MATERIAL AND METHODS

### Cell cultures

Human primary dermal fibroblast cell line C004 (foreskin fibroblast strain #2294, from a 4-year-old donor) was a generous gift from the Laboratory of Molecular and Cell Biology, Istituto Dermopatico dell’Immacolata (IDI)-IRCCS (Rome, Italy), while human primary dermal fibroblast cell line C013 (from a 13-year-old donor) was kindly provided by the Italian Laminopathies Network. Human primary dermal fibroblast cell lines C001 (foreskin fibroblast AG08498, from a 1-year-old donor) and C002 (foreskin fibroblast AG07095, from a 2-year-old donor) were obtained from the Coriell Institute. All human primary cell lines were cultured in DMEM high glucose supplemented with GlutaMAX (Gibco, 10566-016), with further addition of 15% (v/v) FBS (Gibco, 10270106), 100 U/mL penicillin G and 100 μg/mL Streptomycin Sulphate. Mouse myoblast C2C12 cell line (ATCC) was cultured in DMEM high glucose supplemented with 10% FBS. All cell lines were cultured at 37 °C, 5% CO_2_.

### Chromatin fractionation

For 4f-SAMMY-seq, 3 million fibroblasts or 1-2 million myoblasts were detached from the culture plate by 3 min incubation in Trypsin-EDTA solution at 37°C, 5% CO2. After two washes in cold PBS, the cells were resuspended in 600 μL of CSK-Triton buffer (10 mM PIPES pH 6.8, 100 mM NaCl, 1 mM EGTA, 300 mM Sucrose, 3 mM MgCl2, 1 mM PMSF, 1 mM DTT, 0.5% Triton X-100, with protease inhibitors). After 10 min incubation on a wheel at 4°C, soluble proteins and the cytoskeletal structure were separated from the nuclei by centrifugation at 900g for 3 min at 4°C; the supernatant was labelled as S1 fraction. The pellet was then washed with an additional volume of CSK-Triton buffer, resuspended in 100 μL of CSK buffer (10 mM PIPES pH 6.8, 100 mM NaCl, 1 mM EGTA, 300 mM Sucrose, 3 mM MgCl2, 1 mM PMSF, with protease inhibitors) and incubated for 60 min at 37°C with 25 U of RNase–free DNase I (Invitrogen, AM2222). To stop DNA digestion, ammonium sulphate was added in the CSK buffer to a final concentration of 250 mM. After 5 min incubation on ice, the sample was pelleted at 900g for 3 min at 4°C; the supernatant, containing digested chromatin fragments, was labelled as S2 fraction. Afterwards, the pellet was washed with 200 μL of CSK buffer and pelleted at 3000g for 3 min at 4°C, then resuspended in 100 μL of CSK-NaCl buffer (CSK buffer with NaCl final concentration increased to 2 M) and incubated for 10 min on a wheel at 4°C. At the end of the incubation, the sample was centrifuged at 2300g for 3 min at 4°C and the supernatant was labelled as S3 fraction. Finally, after two 10 min washes in 200 μL of CSK-NaCl buffer on a wheel at 4°C followed by centrifugation at 3000g for 3 min at 4°C, the pellet was solubilized in 100 μL of 8 M urea; the final suspension was labelled as S4 fraction. For 50k and 10k 4f-SAMMY-seq, 50,000 and 10,000 fibroblasts were used, respectively. They were processed following the 4f-SAMMY-seq protocol described above but using only 2U of RNase–free DNase I (Invitrogen, AM2222).

For 10kh-SAMMY-seq, 10,000 fibroblasts were processed following the 4f-SAMMY-seq protocol described above but halving the volumes in each step and using 12.5U of RNase– free DNase I (Invitrogen, AM2222); in this way, we achieved a 150-fold increase in the ratio of enzyme units per starting number of cells, while maintaining the DNase concentration unaltered.

### DNA extraction, sonication and sequencing

Fractions S2, S3 and S4 were diluted in TE (10 mM TrisHCl pH 7.5, 1 mM EDTA pH 8.0) to a final volume of 200 μL and then incubated 90 min at 37°C with 6 μL of RNase cocktail (Ambion, AM2286), followed by 150 minutes at 55°C with Proteinase K (Invitrogen, AM2548) to a final concentration of 0.2 μg/μL. Next, DNA was isolated through phenol:chloroform:isoamyl alcohol (Sigma, 77617) extraction, precipitated in 70% ethanol, 0.3 M sodium acetate and 20 μg glycogen overnight at -20°C or 1 hour in dry ice and resuspended in nuclease-free water. S2 from 4f-SAMMY-seq was additionally purified using PCR DNA Purification Kit (Qiagen, 28106) and then DNA fragments in this fraction were separated using AMPure XP paramagnetic beads (Beckman Coulter, A63880) to obtain S2S (< 300bp) and S2L (> 300bp) fractions. In detail, beads were added to the S2 fraction in a 0.95x (v/v) ratio, so to bind fragments larger than 300bp while leaving smaller fragments in solution. Magnetic separation of beads from supernatant allowed the physical separation of larger fragments (on the beads) from shorter ones (in the supernatant). Larger fragments bound on the beads were then washed in 85% ethanol, resuspended in water and magnetically separated from the beads (S2L fraction). Shorter fragments in the supernatant of the first step were bound to beads by adding a further 0.85x (v/v) beads ratio to the suspension; after washing in 85% ethanol and resuspending in water, they were also detached from the beads (S2S fraction). Separation of S2S and S2L from S2 fraction of 10kh-SAMMY-seq was also tested for one sample (C004_r1); since the enrichment profile of the sequenced S2S and S2L fractions was identical, this passage was later avoided (C002_r1, C004_r2). After DNA isolation, S2 (from 10kh-SAMMY-seq), S2L (from 4f-SAMMY-seq), S3 and S4 (from both 10kh- and 4f-SAMMY-seq) fractions were transferred to screw cap microTUBEs (Covaris, 004078) and sonicated in a Covaris M220 focused-ultrasonicator to obtain a smear of DNA fragments peaking at 200bp (settings: water bath 20°C, peak power 30.0, duty factor 20.0, cycles/burst 50; duration: 125 sec for S2 and S2L, 175 sec for S3 and S4). DNA in the fractions was then quantified using Qubit dsDNA HS Assay Kit (Invitrogen, Q32854) and a Qubit 4.0 fluorometer; quality control was performed by run on an Agilent 2100 Bioanalyzer System using the High Sensitivity DNA Kit (Agilent, 5067-4626). Libraries were then created from each fraction using the NEBNext Ultra II DNA Library Prep Kit for Illumina (NEB, E7645L) and the Unique Dual Index NEBNext Multiplex Oligos for Illumina (NEB, E6440S); final qualitative and quantitative controls were performed through an Agilent 2100 Bioanalyzer System and a Qubit 4.0 fluorometer. Libraries with distinct adapter indexes were multiplexed and, after cluster generation on FlowCell, sequenced for 50 bases in paired-end mode on an Illumina NovaSeq 6000 instrument at the IEO Genomic Unit in Milan or for 100 bases in single-end mode on an Illumina NextSeq 2000 instrument at Ospedale Policlinico in Milan. A sequencing depth of at least 24.9 million raw sequencing reads was obtained for each sample.

### Protein analysis

Chromatin fractions were quantified using Qubit Protein Assay Kit (Invitrogen, Q33212) and a Qubit 4.0 fluorometer. Equal protein amounts from each fraction were run on 4-12% bis-tris plus acrylamide gels (Invitrogen, NW04122) and then immunoblotted. Anti-tubulin alpha (Sigma T5168, mouse 1:2000), anti-H3 (Abcam ab1791, rabbit 1:4000 or Invitrogen #MA3-049, mouse 1:1000), anti-Lamin A/C (Santa Cruz sc-6215, goat 1:1000) and anti-Lamin B (Santa Cruz sc-6216, goat 1:2000) were diluted in 5% (w/v) milk in PBST (0.1% Tween in PBS) and used as primary antibodies. Secondary HRP-conjugated anti-mouse (Sigma, A9044), anti-rabbit (Sigma, A9169) and anti-goat (Sigma, A5420) antibodies were then revealed through SuperSignal West Dura chemiluminescence kit (Thermo Scientific, 34076) and signals acquired in an iBright FL1500 Imaging System.

### Public datasets

We used publicly available 3f-SAMMY-seq datasets from our previously published article on SAMMY-seq (GEO dataset GSE118633).

We collected publicly available human ChIP-seq and DNase-seq datasets from the following sources: Lamin A/C ChIP-seq from (17) (SRA datasets IDs SRR605493, SRR605494, SRR605495 and SRR605496); Lamin B1 ChIP-seq from (20); ChIP-seq for H3K27ac, H3K36me3, H3K4me1, H3K4me3 and H3K9me3 from Roadmap Epigenomics (primary human foreskin fibroblast from newborn, sample E055)(21); ChIP-seq for H3K27me3 from our previously published data (16)(Sample CTR002); DNAse-seq from Roadmap epigenomics (primary human foreskin fibroblast from newborn, sample E055)(22).

We collected publicly available mouse ChIP-seq from the following sources: H3K27me3, H3K4me1, H3K4me3 from (23), H3K36me3 from the Mouse Encode consortium (GEO sample GSM918417)(24) and H3K9me3 from (25).

We also downloaded from the Roadmap Epigenomics portal (https://egg2.wustl.edu/roadmap/data/byDataType/rna/expression/57epigenomes.RPKM.pc.gz) the RNA-seq based gene expression profiles (RPKM) of protein coding genes for human foreskin primary fibroblasts (sample E055). Genes coordinates were also downloaded from the same Roadmap Epigenomics portal (Ensembl_v65.Gencode.V10.ENSG.gene.info). Genes coordinates have been converted from hg19 to hg38 reference genome assemblies using UCSC tool hgLiftOver, setting “Minimum ratio of bases that must remap” to 1 and “Allow multiple output regions” to disabled.

For human fibroblasts we downloaded the publicly available Hi-C pre-processed aligned, filtered and binned read counts in .mcool file format from the 4DN portal (26)(HFF-hTERT, dilution protocol, HindIII - sample ID 4DNFIMDOXUT8: raw multiple-resolution contact matrix). Where needed reads counts were aggregated to address a limitation of the cooler pipeline used by 4DN consortium as per their suggested guidelines (https://data.4dnucleome.org/files-processed/4DNFIJTOIGOI/).

For mouse myoblasts (C2C12 cells) we downloaded Hi-C raw reads (FASTQ) from GEO (dataset GSE104427) from (27).

### High-throughput sequencing reads alignment and filtering

High-throughput sequencing reads were trimmed using Trimmomatic (v0.39) (28) using the following parameters for SAMMY-seq and ChIP-seq: 2 for seed_mismatch, 30 for palindrome_threshold, 10 for simple_threshold, 3 for leading, 3 for trailing and 4:15 for sliding window. The sequence minimum length threshold of 35 was applied to all datasets. We used the Trimmomatic “TruSeq3-SE.fa” (for single-end) and “TruSeq3-PE-2.fa” (for paired-end) as clip files. After trimming, reads were aligned using BWA (v0.7.17-r1188) (29) setting –k parameter as 2 and using as reference genome the UCSC hg38 genome or mm9 genome (only canonical chromosomes were used) for human and mouse datasets, respectively. The output aligned reads were saved in BAM file format. We uniformly aligned and used only a single read per DNA fragment for both single-end and paired-end sequencing samples. The PCR duplicates were marked with Picard (v2.22; https://github.com/broadinstitute/picard) MarkDuplicates option, then filtered using Samtools (v1.9) (30). In addition, we filtered all the reads with mapping quality lower than 1. Each sequencing lane was analysed separately and then merged at the end of the process. For C002-rep1 and C004-rep2 two sequencing runs from the same library were produced and merged after alignment and filtering.

The coverage estimation (as reported in Supplementary Figure 1e) has been performed using the Samtools (v1.16.1) “coverage” command.

For the mouse C2C12 Hi-C dataset, raw reads (FASTQ) were aligned, filtered, binned and the count matrix was calculated using the NextFlow implementation of HiC-Pro (31): namely the “hic” pipeline (version 2.1.0) from the nf-core repository (32). The reference genome used for the alignment was the mouse genome version mm9 downloaded from UCSC. The mapping quality minimum (parameter –min_mapq) was set as 10.

For both human and mouse Hi-C datasets, the .mcool files with raw counts were processed with cooler (version 0.8.3)(33) for matrix balancing normalization with iterative bias correction (ICE)(34). Normalized contact matrices at specific resolutions (binsize: 250000 and 50000) were exported in text format with the “cooler dump” function for each chromosome, then processed for subsequent analyses as described in specific sections below.

### High-throughput sequencing reads analyses

To compute reads distribution profile (genomic tracks), we used DeepTools (v3.4.3) (35) bamCoverage function. For this analysis the genome was binned at 50bp, the reads extended up to 250bp and RPKM normalization method was used. We considered a genome size of 2,701,495,761bp (value suggested in the DeepTools manual https://deeptools.readthedocs.io/en/latest/content/feature/effectiveGenomeSize.html) and we excluded regions known to be problematic in terms of sequencing reads coverage using the blacklist from the ENCODE portal (https://www.encodeproject.org/files/ENCFF356LFX/).

To compute the genomic tracks for ChIP-seq IP over INPUT enrichment profiles (log2 normalized ratio) or for relative comparisons (relative enrichment, i.e. log2 normalized ratio) between SAMMY-seq fractions, we used the SPP R package (v1.16.0) (36) and R statistical environment (v3.5.2). The reads were imported from the BAM files using the “read.bam.tags” function, then filtered using “remove.local.tag.anomalies” and finally the comparisons were performed using the function “get.smoothed.enrichment.mle” setting “tag.shift = 0” and “background.density.scaling = TRUE”.

To compute correlations between genomic tracks, we used R (v3.5.2) base function “cor” with “method = Spearman”. The genomic tracks were imported in R using the rtracklayer (v1.42.2) (37) library. Then the files were binned at 50kb using the function “tileGenome” and the correlation was computed per chromosome. The correlation values obtained for each chromosome were then summarized in one value describing the genome-wide sample correlations through a weighted mean, where the weight of each chromosome corresponds to its length.

To compute the average profile over a set of genomic regions of interest (i.e. the meta-profile analysis), we used DeepTools (v3.4.3)(35) “computeMatrix” command. For gene centred meta-profiles (see Supplementary Figure 2b) we used as regions of interest the protein coding genes annotated by RefSeq database (genome version hg38) that are not overlapping to each other considering a window 2.25kb upstream and 0.25kb downstream the TSS. The binning value was set to 10bp, the body region was rescaled to 3,000bp and the flanking regions included up to 2Kb. In addition, the “skipZeros” option was added to remove regions with no coverage. The genes were separated by quartile of expression using the reference expression profiles (E055) from Roadmap Epigenomics after separating the genes with 0 RPKM. The meta-profile plots were then drawn using the “plotProfile” tool of DeepTools using as input the previously created matrix.

To quantify the differences in meta-profiles around TSS, the paired effect size (Cohen’s *d*) (38,39) between reference genomic positions relative to TSS (Supplementary Figure 2c) was calculated. Namely, we computed the mean reads count for each replica in 500bp windows around TSSs and in 500bp windows 2Kb upstream of the TSSs. Only protein-coding genes without overlaps with other transcripts in the regions considered for the effect size calculation (*i.e.*, 2.25Kb upstream and 0.25 downstream the TSS) were kept.

For meta-profiles centred over genomic regions marked by peaks enriched for specific chromatin marks (see Figure 1c and Supplementary Figure 3), we analysed ChIP-seq data for histone marks with parameters adapted to the type of enrichment profiles: either sharp (H3K4me1, H3K4me3, H3K27ac), broad (H3K27me3, H3K9me3) or broad and associated to gene bodies (H3K36me3). Namely, for sharp histone marks (H3K4me1, H3K4me3, H3K27ac) we called enrichment peaks with MACS (v2.2.9.1) (40) using a broad-cutoff of 0.1. We then merged close peaks for H3K4me1 and H3K27ac (a distance threshold of 500bp and 400bp was used, respectively), using BEDTools (v2.29.1) (41). Finally, for every sharp histone mark, peaks that were not overlapping to each other considering an extended window of 1Kb (+/- 500bp around the peak) were considered as regions of interest. DeepTools (v3.5.1)(35) “computeMatrix” command was used to extract the matrix of reads distribution profiles of SAMMY-seq fractions on the regions, setting the binning at 50bp, rescaling the domain body region to 1Kb and including flanking regions of 1Kb. In addition, the “skipZeros” option was added to remove regions with no coverage. The meta-profile plots were then drawn using the “plotProfile” tool from DeepTools using as input the previously created matrix. The number of regions displayed for H3K4me1, H3K4me3 and H3K27ac is 117593, 50299 and 23404, and their average length is 1380bp, 1089bp and 875bp, respectively. For H3K27me3, we called enrichment peaks with MACS, but we used a broad-cutoff of 0.05 and then used BEDTools to merge peaks that were less than 1Kb apart. Finally, peaks that were not overlapping to each other considering an extended window of 3Kb (+/-1500bp flanking the peak) were considered as regions of interest. The DeepTools “computeMatrix” was used as described above but with binning at 50bp, rescaling the body region to 3Kb and including flanking regions of 3Kb, and the meta-profile drawn using the “plotProfile” function. The number of regions used for the meta-profile is 22418, with an average length of 2064bp. For broad histone mark H3K9me3, we called enrichment peaks with EDD (v1.1.19)(42), using a bin-size of 40Kb and a gap-penalty of 25. Peaks that were not overlapping to each other considering an extended window of 1.5Mb were considered as regions of interest. The DeepTools “computeMatrix” command was used as described above but with binning at 30Kb, rescaling the body region to 1.5Mb and including flanking regions of 1.5Mb, and the meta-profile drawn using the “plotProfile” function. The number of regions used for the meta-profile is 187, with an average length of 1581818bp. For H3K36me3, i.e. a histone mark with broad enrichment profile but usually associated to the body of transcribed genes, we called enrichment peaks with MACS using a broad-cutoff of 0.1. We then used NCBI RefSeq annotations for hg38 genome build (https://hgdownload.soe.ucsc.edu/goldenPath/hg38/bigZips/genes/hg38.ncbiRefSeq.gtf.gz) to associate each peak with its overlapping protein coding gene body. We also removed genes close to each other if they had a H3K26me3 peak in common, thus with potential ambiguous peak to gene association. Finally, we merged peaks in case more than one is associated to an individual gene body and the resulting peaks that were not overlapping considering an extended window of 10Kb were considered as regions of interest. DeepTools (v3.5.1) “computeMatrix” command was used as described above but with binning at 50bp, rescaling the body region to 60Kb and including flanking regions of 10Kb. The number of regions used for the meta-profile is 5990, with an average length of 62821bp. For the “oriented” version of H3K36me3 meta-profile plot, we oriented the regions in line with the strand of transcription of the corresponding gene body.

**Figure 1.**
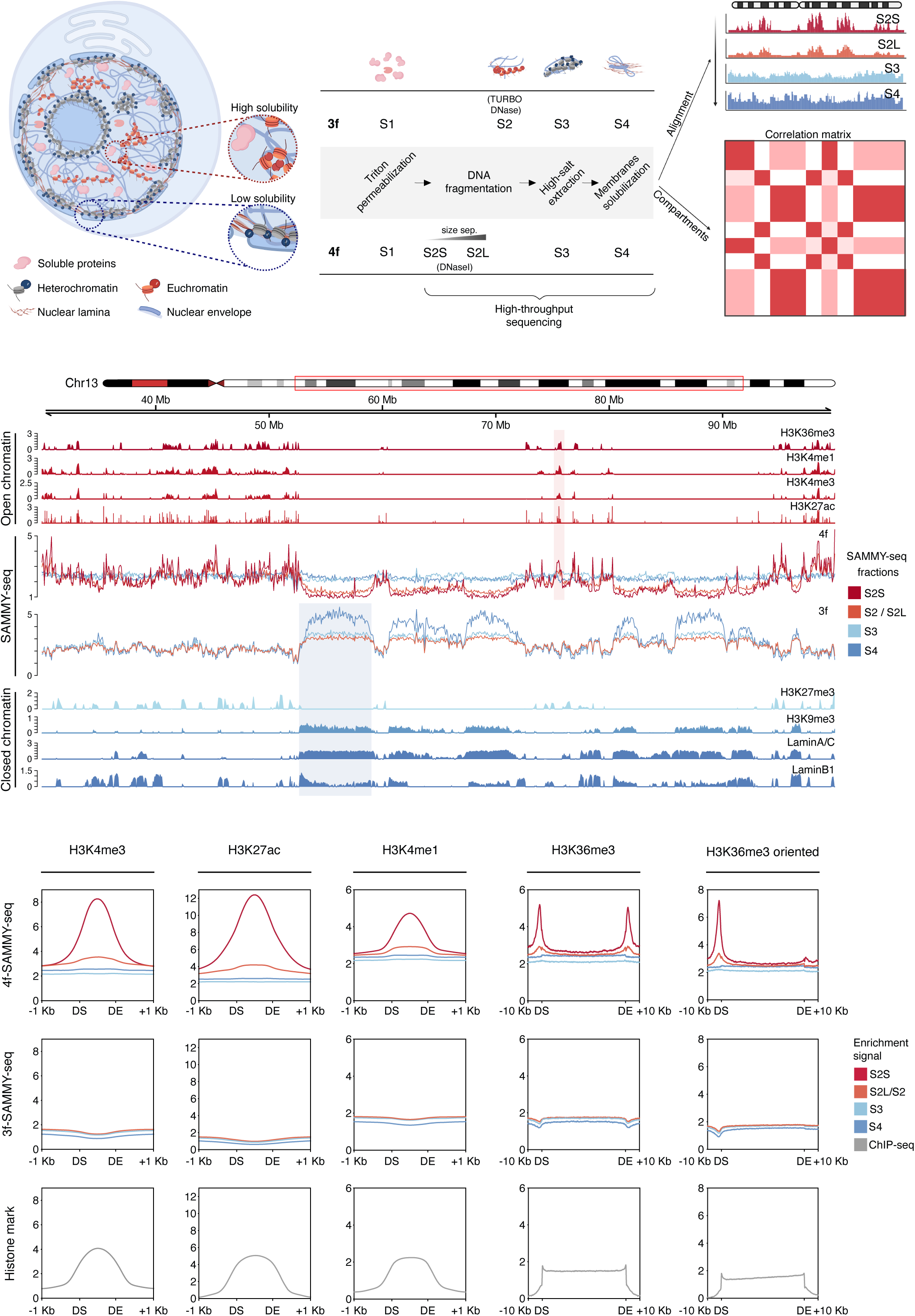
4f-SAMMY-seq maps both euchromatin and heterochromatin with high resolution. **a)** Schematic illustration of the 3f-SAMMY-seq vs 4f-SAMMY-seq protocols and output analysis results. From left to right, high (euchromatin) and low solubility (heterochromatin) domains correspond to genomic portions with preferential segregation in different subnuclear regions. In both “3f” and “4f” SAMMY-seq protocols, a sequential extraction of chromatin fractions (numbered from S1 to S4) results in a separation of euchromatic and heterochromatic regions that are then mapped to their genomic coordinates by high-throughput sequencing (applied to fractions from S2 through S4). The novel 4f-SAMMY-seq has a lighter digestion obtained with DNase I replacing TURBO DNase. Moreover, the S2 fraction is size separated into short (S2S) and long (S2L) fragments before sequencing. Specific data analysis procedures allow reconstructing chromatin domains compartmentalization in the 3D nuclear space. **b)** Representative genomic region (chr13:30,000,000-100,000,000) showing genomic tracks for chromatin marks in human foreskin fibroblasts. From top to bottom: open chromatin marks ChIP-seq enrichment profiles for H3K36me3, H3K4me1, H3K4me3, H3K27ac; reads distribution profiles for individual fractions of a representative replicate of 4f-SAMMY-seq (C004_r2) and 3f-SAMMY-seq (C004_r1); closed chromatin marks ChIP-seq enrichment profiles for H3K27me3, H3K9me3, Lamin A/C, Lamin B1. The shaded areas mark two examples of regions showing enrichment for closed (blue) or open (red) chromatin marks. **c)** Reads distribution meta-profiles for individual “4f” or “3f” SAMMY-seq fractions in the first and second row of plots, respectively. The bottom row of plots reports the reads distribution profile of the corresponding histone mark IP samples from ChIP-seq experiments. The average reads distribution profiles are computed over chromatin domains marked by enrichment peaks of specific histone marks, indicated on the top of each column. The domain start (DS) and domain end (DE) are indicated on the x-axis along with flanking regions coordinates. For H3K36me3 mark, we reported also the meta-profile obtained by orienting the domains according to the corresponding gene’s transcribed strand. Meta-profiles for additional replicates are reported in Supplementary Figure 3a.

To study distribution of gene expression levels across the compartments (see Supplementary Figure 6a), for each gene we considered a window of +/-500bp around the TSS. These windows were intersected with compartments coordinates using BEDTools (v2.29). 2928 genes were discarded because their coordinates are not present in the compartment classification, while 22 genes were discarded because they overlap with more than one compartment. Of the remaining 16656 genes, 2721 (with RPKM = 0) were classified as “Not detected”; on the RPKM of the others, we applied R function “quantile” to classify genes under 25% in “Low”, between 25% and 50% in “Moderate”, between 50% and 75% in “High” and over 75% in “Very high”.

### Chromatin states analysis

We used chromHMM (v1.24)(43) to call chromatin states with a 15 states model for both human fibroblasts and mouse C2C12 cells. In line with the approach previously adopted by the Roadmap Epigenomics consortium (21), for each of these cell types we used a set of histone marks ChIP-seq data including: H3K4me1, H3K4me3, H3K36me3, H3K27me3, H3K9me3. To obtain the chromatin segmentation model, we initially used the ‘BinarizeBed’ function to binarize the input data based on the observed count of sequencing reads according to a Poisson distribution with specified parameters: bin size of 200bp and a p-value threshold of 0.0001. The function discretizes each histone mark ChIP-seq bin into two levels, 1 indicating enrichment and 0 indicating no enrichment. Afterwards, we train a multivariate Hidden Markov Model (HMM) of 15 states using the default parameters of the function ‘LearnModel’. Thus, we manually curated the labelling of chromatin states in order to maximize comparability with the consensus definition of chromatin states previously described in (21). The output 15 chromatin states from each analysis were merged in case two states had very similar redundant emission state probability profiles. As a result of this manual curation, the final set of chromatin states is 14 for the human fibroblast dataset and 13 for the mouse myoblast dataset.

The chromatin marks enrichment matrix associated with chromHMM chromatin states for human fibroblasts (Supplementary Figure 6b) was computed as the average ChIP-seq IP over INPUT enrichment (computed with SPP package as described above) for each chromatin mark and state. Given the different range of values, the colour gradient in the heatmap has been rescaled, for each chromatin mark, to the maximum enrichment for that mark. For SAMMY-seq fractions, the average enrichment is computed from RPKM of reads distribution profiles and rescaled as a z-score to allow comparison of different SAMMY-seq fractions with distinct enrichment ranges.

### Genomic tracks visualization

The visualization of genomic tracks was performed with Gviz R library (v1.26.5) (44). The track profile was calculated using the function “DataTrack” (the input file was imported using the function “import” of the rtracklayer library) and plot using the function “plotTracks” setting the value “window = 1000”. Line plots were drawn setting the parameter type as ‘a’ and overlayed using the function “OverlayTrack”; instead, mountain plots were obtained setting the parameter type as “polygon”. Extra elements of these plots, such as chromosome ideogram (on top) and genome axis, were plotted respectively using the functions “IdeogramTrack” and “GenomeAxisTrack”. The analysis was performed identically for all the datasets except for the H3K27ac tracks, where the “window” parameter was set to 10,000 to improve the genomic tracks visualization.

### Chromatin compartments analysis

We used the Hi-C and SAMMY-seq pairwise correlation matrices of normalized contacts and reads distribution profiles, respectively, binned at 250 kb (Figure 2a). Bins with null contacts or coverage were removed from both matrices.

**Figure 2.**
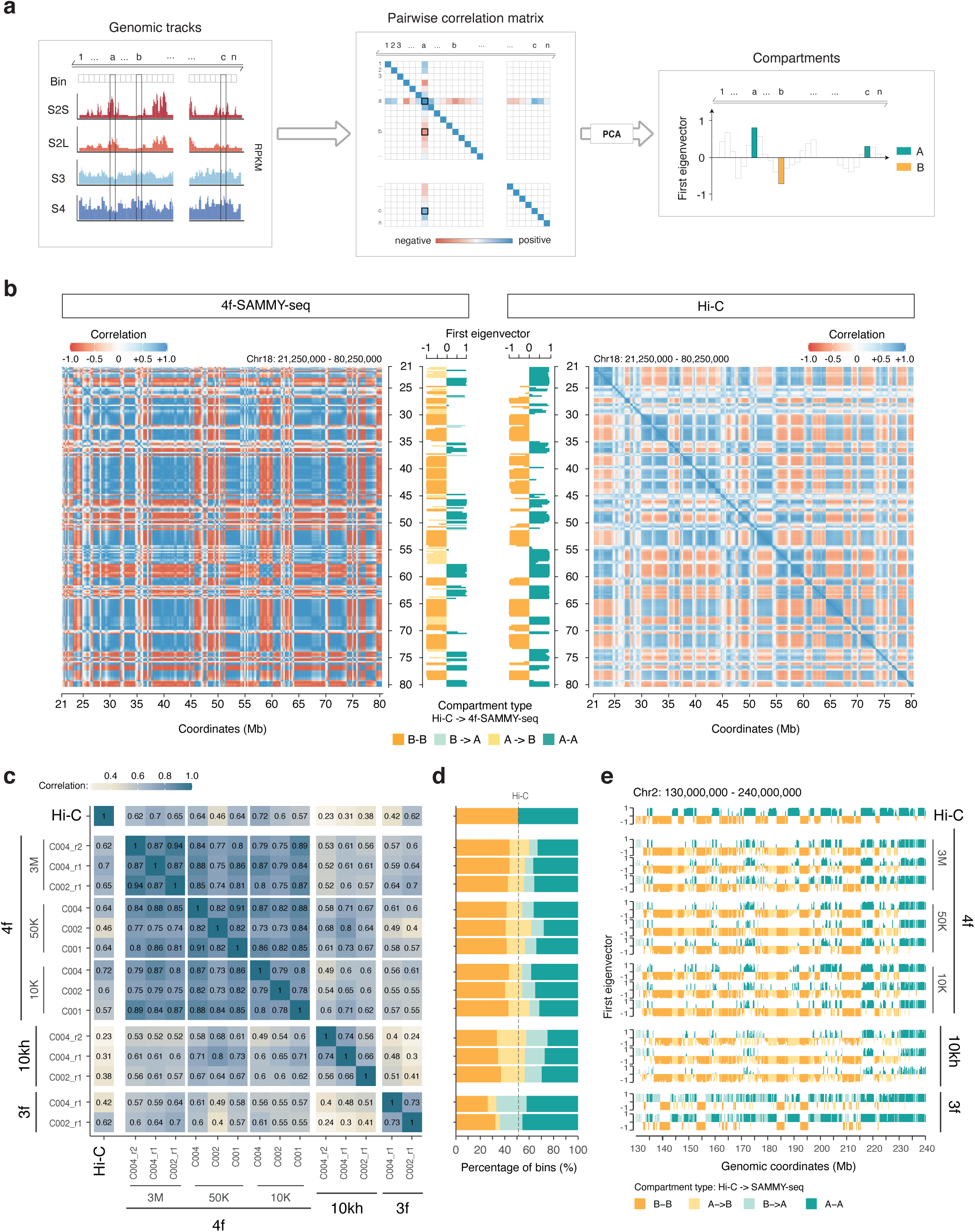
4f-SAMMY-seq detects chromatin domains segregation in compartments. **a)** Schematic illustration of the data analysis workflow to reconstruct chromatin compartments from SAMMY-seq data. For the “A” and “B” compartments identification, starting from read distribution profiles for individual chromatin fractions. The Pearson correlation is computed chromosome-wise between the vectors of reads coverage across fractions for each pair of genomic bins. After performing a principal component analysis (PCA), the first component (corresponding to the first eigenvector of the matrix) is used to discriminate active “A” compartment (positive values) and inactive “B” compartment (negative values) (see also Methods). **b)** Pairwise correlation matrices of read distribution profiles for 4f-SAMMY-seq fractions (C002_r1, left) or distance normalized (observed over expected) Hi-C contact profiles (right) on a representative genomic region (chr18:21,250,000-80,250,000) computed at 250Kb genomic bins resolution. On the side of each matrix the respective first eigenvector is reported and coloured to mark the position of active (”A” compartment) and inactive (”B” compartment) regions. Concordant domain classification in Hi-C and 4f-SAMMY-seq are marked in orange (”A-A”) for active regions and green (”B-B”) for inactive regions. In the 4f-SAMMY-seq eigenvector only, we marked differently the regions with a discordant compartment classification: a lighter green is used for regions classified as B in Hi-C and A in 4f-SAMMY-seq (”B->A” label), a lighter orange is used for the opposite case (”A->B” label). **c)** Genome-wide pairwise Pearson correlation of chromatin compartments eigenvectors defined by Hi-C and SAMMY-seq protocol variants “3f”, “10kh” and “4f” starting from 3 million (3M), 50,000 (50K) or 10,000 (10K) cells with individual replicates reported. **d)** For each sample of panel c, the stacked barplot shows the relative percentage distribution of genomic bins associated with concordant (”A-A” or “B-B”) and discordant (”A->B” or “B->A”) compartment classification with respect to Hi-C. The same colouring and naming convention of panel b was adopted. The samples order is the same as in panel c. **e)** Chromatin compartments eigenvectors for a representative genomic region (chr2:130,000,000-240,000,000). The samples order is the same as in panel c. The eigenvectors are coloured according to the same convention adopted in panel b.

Namely, for the Hi-C protocol, we first: i) balanced the raw contact matrix (cooler balance -- cis-only, version 0.8.3), ii) loaded the ICE balanced chromosome-wise contact matrix (cooler dump, version 0.8.3), and iii) normalized it by dividing each observed pairwise contact by the mean of the contacts at the same distance (*i.e.*, the expected); secondly, we computed the correlation between all pairs of bins. This step required the calculation of *n*x*n* correlations of *n*-dimensional vectors of normalized contacts, where *n* is the number of bins. For the SAMMY-seq protocols, we loaded the bin enrichment tracks (RPKM) of each fraction and computed the correlation between pairs of bins. Here, the correlation was computed between *m*-dimensional vectors of reads distribution profiles, where m was the number of fractions in the specific protocol (*i.e.*, m=3 for the 3f and 10k protocols, m=4 for the 4f protocol).

For each input correlation matrix, the first eigenvector was obtained through principal component analysis decomposition in R statistical environment (prcomp, stats package, center=FALSE and scale=TRUE, rotation component of the returned object). The sign of the first eigenvector was defined using gene density: the group of bins with the highest gene density was marked as “A” compartment (positive sign), and the group with the lowest gene density was marked as “B” (negative sign). Chromosome eigenvector values were divided by the absolute maximum value for visualisation purposes. All the analyses were made using R (version 3.6.0).

### Sub-compartments analysis

We partially reimplemented CALDER algorithm (version 1.0, 2020-09-01) to accommodate the different format of SAMMY-seq data, maintaining the primary set of CALDER core functions (remove_blank_cols, fast_cor, generate_compartments_bed, HighResolution2Low_k_rectangle, get_PCs, bisecting_kmeans, project_to_major_axis, get_best_reorder, get_cluser_levels), their default parameters and the steps intended in the original article (8).

As inputs, we used the balanced chromosome-wise contact matrix (cooler dump, version 0.8.3)(33) and the pairwise distance matrix binned at 50 kb for Hi-C and SAMMY-seq protocols, respectively (Figure 5a). In particular, the pairwise distance matrix was computed using for each pair of bins the Euclidean distance of the m-dimension vectors, where m was the number of fractions in the specific protocol (*i.e.*, 4 for the 4f-SAMMY-seq protocol). Bins with null contacts or reads coverage were removed from both matrices.

Briefly, for each input matrix, the algorithm computed the pairwise correlation matrix and identified the sub-compartments boundaries; computed the binary trend matrix and its decomposition using ten principal components; iteratively clustered the domains to obtain their hierarchy; sorted the domains hierarchy based on the projection of the first two components on the lowess interpolation; divided them into eight groups (form the most closed, *i.e.*, B.2.2 to the most opened, i.e., A.1.1). For more details on the original CALDER procedure and the interpretation of sub-compartments refer to (8).

In the analysis of chromatin marks enrichment associated with sub-compartments (Figure 5b, Supplementary Figures 6e and 7c): i) we computed the average ChIP-seq IP over INPUT enrichment for each bin (SPP package, as described above); ii) a z-score normalization by chromosome (i.e., we subtracted the mean and divided it by the standard error) was applied, and iii) we calculated the mean of the z-scores (i.e., the normalized ChIP-seq IP over INPUT enrichment) by sub-compartment. This relative scaling normalization step was applied to improve comparability across different chromosomes.

## RESULTS

### High resolution mapping of open chromatin regions by solubility

We previously proposed the SAMMY-seq method based on the sequential biochemical purification and sequencing of three distinct chromatin fractions, hereafter called “3f” SAMMY-seq (16). These fractions were respectively isolated after partial digestion with an endonuclease (TURBO DNase, fraction S2), extraction with high salt concentration to dissolve ionic bonds (fraction S3), and treatment with urea buffer to dissolve the remaining protein and membrane pellet (fraction S4) (Figure 1a and Supplementary Figure 1a). The latest fractions of 3f-SAMMY-seq (S3 and S4) are enriched for the compacted and less soluble heterochromatin domains (Figure 1b, Supplementary Figure 1a and 2a), whereas none of the chromatin fractions could precisely reflect individual sharp peaks of chromatin accessibility, such as those usually found at actively transcribed genes and regulatory elements. To improve the resolution in characterizing the accessible euchromatin, we operated two crucial changes in the protocol. First, we introduced a lighter DNase (DNaseI) digestion in preparation for S2 fraction isolation (Figure 1a and Supplementary Figure 1b, see Methods for details). Second, we introduced a size selection step on the S2 fraction DNA, to further separate more digested small-size DNA fragments (S2-Short, S2S, <300bp) from less digested large-size DNA fragments (S2-Long, S2L) (Figure 1a and Supplementary Figure 1c). The resulting four chromatin fractions were sequenced, thus we named this the “4f” SAMMY-seq protocol.

We applied 4f-SAMMY-seq on human primary fibroblasts and compared the results to 3f-SAMMY-seq data on the same cells. We confirm good and comparable results in terms of quality controls on the proteins associated with each fraction (Supplementary Figure 1a) as well as on the sequencing reads, achieving good coverage and reproducibility even with moderate sequencing depth (minimum of 25 million reads per fraction) (Supplementary Table 1 and Supplementary Figure 1d-f). The S2S and S2L fractions are able to precisely recapitulate the location of histone marks associated with active chromatin (H3K36me3, H3K4me1, H3K4me3 and H3K27ac) (Figure 1b-c, and Supplementary Figures 2a-b and 3a). When examining 3f-SAMMY-seq vs 4f-SAMMY-seq fractions only the latter one has an evident enrichment profile over domains marked by histone marks of active chromatin (Figure 1c and Supplementary Figure 3a). Moreover, the most accessible S2S fraction achieves a level of detail comparable to DNase-seq (45) in recapitulating the accessibility profiles around the transcription start site (TSS) of annotated genes, also in their association with expression level (Supplementary Figure 2b). We also quantified the different sensitivity in detecting a TSS profile enrichment in relation to expression quantile with an effect size analysis (Supplementary Figure 2c). This analysis showed that 4f-SAMMY-seq performs better than 3f-SAMMY-seq in terms of reads enrichment at TSS across all expression levels.

These results confirm that the novel 4f-SAMMY-seq is overcoming a crucial limitation of 3f-SAMMY-seq by achieving a detailed and quantitative map of active chromatin regions. Overall, these data support our hypothesis that the combination of controlled digestion with fragments size separation allows a more precise characterization of open chromatin regions.

### Active and inactive chromatin mapping on a small number of cells

We asked if the novel 4f-SAMMY-seq is also able to identify lamina-associated constitutive heterochromatin, on which 3f-SAMMY-seq showed high sensitivity (16,46). In this respect, the relative sequencing reads enrichment in closed (S3) versus open (S2S) chromatin fractions in 4f-SAMMY-seq shows comparable or better results with respect to 3f-SAMMY-seq (Supplementary Figure 4a,b). We noted that this result is driven by S2S reads depletion over heterochromatin regions; in fact S3 and S4 fractions from 4f-SAMMY-seq are not clearly enriched in closed chromatin, as they were instead in 3f-SAMMY-seq (Figure 1b and Supplementary Figure 3b). The lack of heterochromatin enrichment in insoluble S3 and S4 fractions of 4f-SAMMY-seq may be expected given the lighter endonuclease digestion, possibly leaving a mixture of open and closed chromatin in the latest fractions (Supplementary Figure 1b). To confirm this hypothesis, we performed the “4f” chromatin fractionation starting from 10,000 cells and scaling up the ratio of enzyme units per starting number of cells (see methods), so as to achieve a higher level of DNA digestion without changing the endonuclease enzyme. We called this protocol 10kh-SAMMY-seq, whereas previous experiments were generally performed starting from 3 million cells and marked as “3M” in the figures to distinguish from other protocol variants (see also Methods). In this condition, we confirm that the S3 fraction is clearly enriched in heterochromatic regions (Supplementary Figures 1d-e, 2a and 4c). Of note, in the 10kh-SAMMY-seq protocol the S4 is not as informative, possibly due to the limited amount of DNA left in the last fraction when starting from 10,000 cells (Supplementary Figure 2a).

We then tested the possibility of performing 4f-SAMMY-seq starting from as little as 50,000 (4f-50k) or 10,000 (4f-10k) cells while scaling down the DNAse enzyme units (see methods). In these conditions, we can recapitulate the results obtained with 3 million cells (3M) (Supplementary Figures 1d-e, 2a and 4c).

These results confirm the applicability of 4f-SAMMY-seq on as little as 10,000 cells and the possibility of modulating the enrichment for heterochromatin in the less soluble fractions by tuning the endonuclease partial digestion step.

### 4f-SAMMY-seq detection of chromatin compartments

Chromatin 3D compartmentalization is commonly identified based on the long-range similarity of contact profiles measured by Hi-C for each pair of genomic loci. We hypothesized that chromatin domains spatially located in the same 3D nuclear neighbourhood would also be characterized by a similarity of biochemical properties. Indeed, chromatin compartments are characterized by a specific chromatin state, thus exposed to the same biochemical milieu determined by the local concentration of DNA, RNA, associated proteins and epigenetic modifications. All of the SAMMY-seq protocol variants discussed above are based on the biochemical fractionation of chromatin, thus providing a readout on biochemical properties of chromatin in terms of accessibility and solubility.

We compared each pair of genomic regions by computing their “biochemical similarity” as the correlation in the sequencing reads coverage across SAMMY-seq fractions (see Methods and Figure 2a). The resulting 2D matrix is similar to the pairwise correlation matrix of normalized Hi-C data on human fibroblasts (26) (Figure 2b). The eigenvector decomposition of the matrices confirms that the linear segmentation in “A” and “B” compartments is mostly concordant between Hi-C and SAMMY-seq protocols (Figure 2c-e). Among all of the SAMMY-seq protocols, the compartments eigenvectors derived from 4f-SAMMY-seq are consistently showing the highest correlation with the Hi-C eigenvector (Figure 2c), and the highest median Jaccard Index for concordance of compartments across replicates equal to 0.73, 0.62 and 0.78 for 3f, 10kh and 4f-SAMMY-seq (3M), respectively (Figure 2d). In addition, 4f-SAMMY-seq shows the highest performances (eigenvector correlation and reproducibility), also when starting from as little as 50k or 10k cells, with median Jaccard Index across replicates equal to 0.76 and 0.75, respectively (Figure 2d, e). We further confirmed these results in a completely independent set of data for C2C12 mouse myoblast cells, thus covering a different organism and cell type. In this dataset, we observe as well high correlation with Hi-C eigenvectors and concordance of compartments classification (Supplementary Figure 5a).

### 4f-SAMMY-seq based chromatin compartments consistently recapitulate chromatin states

We then analysed in details the differences between 4f-SAMMY-seq-based and HiC-based compartmentalization in terms of transcriptional activity and epigenomics features. Specifically, we focus on the discordant compartments, that were labelled as “A->B” if called as active by Hi-C and inactive by SAMMY-seq, or vice versa for those labelled “B->A” (Figure 2b, d, e). We examined the transcriptional activity in the Hi-C vs SAMMY-seq-based compartments for human fibroblasts. Both techniques showed similar pattern of compartments distribution across distinct gene expression levels, with A compartments assignment directly correlated with transcription. However, 4f-SAMMY-seq showed a tendency to classify more genes in the A compartment (Supplementary Figure 6a).

We further examined chromatin states in genomic regions with discordant compartment classification. We used a combination of proprietary and public reference datasets from Roadmap Epigenomics (21) to define chromatin states (see Methods and Supplementary Figure 6b). An example of the resulting information is provided in Figure 3a, where for each bin the compartment classification and the chromatin states composition are depicted. Of note, in this analysis, dividing the genome in multiple bins of 250 kb, multiple chromatin states can coexist within individual bins. The most prominent and recurrent difference is on chromatin states associated with Polycomb regulation (Figure 3b), including monovalent repressive Polycomb (ReprPC, ReprPCWk) and bivalent chromatin states (EnhBiv, BivFlnk or TssBiv). This was confirmed by results obtained in both human fibroblasts (3M, 50k and 10k cells datasets) and mouse C2C12 datasets (Figure 3a-b, Supplementary Figure 5b-c, 6c, 7a-b).

**Figure 3.**
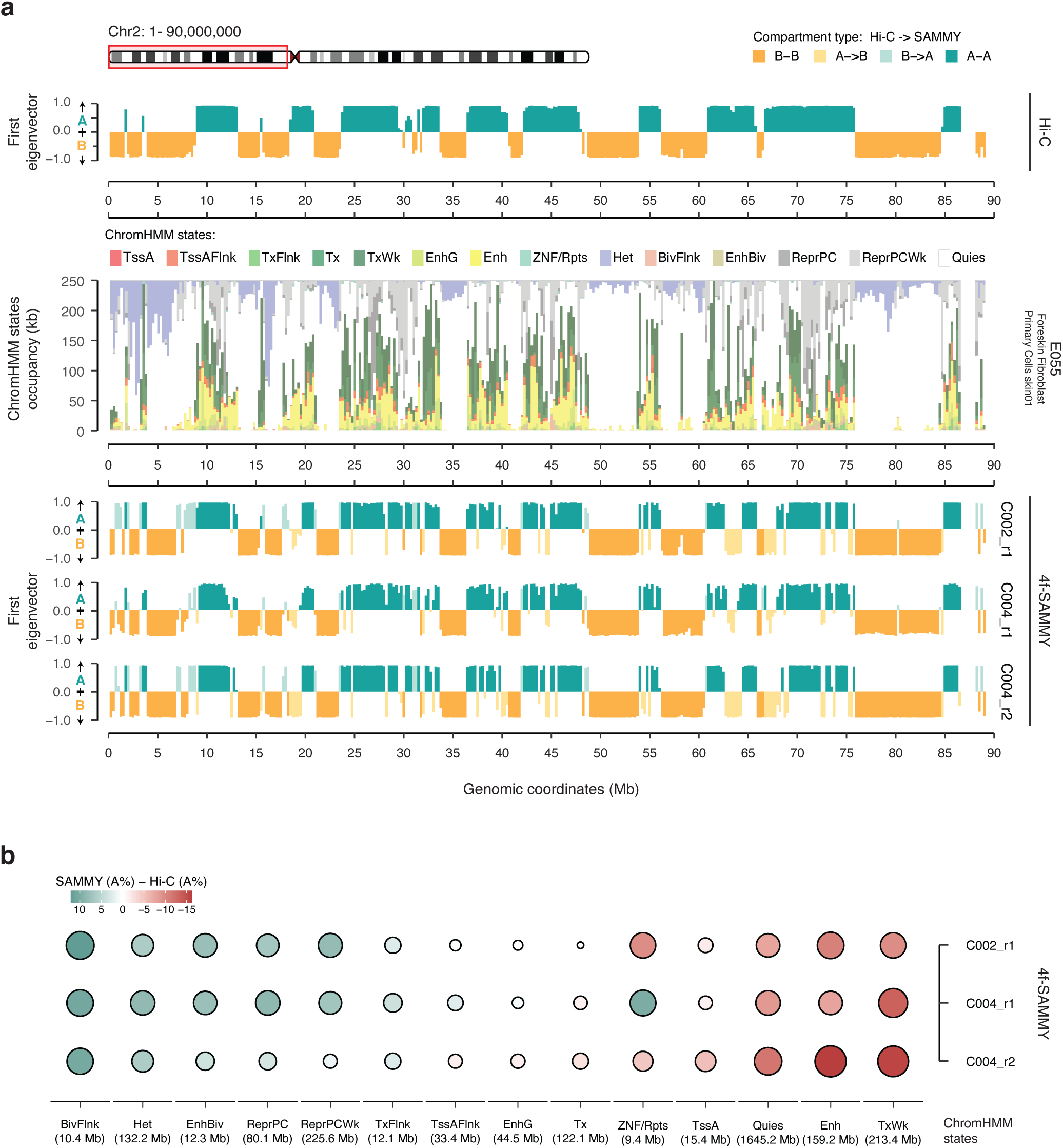
4f-SAMMY-seq based compartments provide a detailed characterization of chromatin transcription and epigenetic status. **a)** Classification of “A” and “B” compartments from Hi-C and 4f-SAMMY-seq and chromHMM chromatin states in human fibroblasts (see Methods) for a representative region (chr2:1-90,000,000). The eigenvector tracks (green and orange tracks) show compartments computed from Hi-C data (top row) and individual 4f-SAMMY-seq replicates (bottom three rows) coloured according to the same convention adopted in Figure 2b for concordant (”A-A” or “B-B”) and discordant (”A->B” or “B->A”) compartments classification. The stacked barplot in the middle row summarizes the chromatin states associated with each genomic bin: see colour legends for states labels and Supplementary Figure 4a for associated chromatin marks signatures. **b)** Relative occupancy of 4f-SAMMY-seq vs Hi-C based compartments for each chromatin state, computed as the difference in “A” compartment percentage. For each chromatin state, positive values (green gradient) indicate a higher percentage of “A” compartment in 4f-SAMMY-seq-based classification, whereas negative values (red gradient) indicate a higher percentage of “A” compartment based on Hi-C classification (i.e. higher “B” percentage based on 4f-SAMMY-seq). The size of each dot is proportional to the absolute value in the percentage difference. Chromatin states are ordered from left to right based on the average percentage difference between 4f-SAMMY-seq replicates (average between the three replicates) and Hi-C.

In our analyses, Hi-C classifies Polycomb-regulated regions prevalently in A compartment with some exceptions assigned to the inactive “B” compartment (Supplementary Figure 6c). On the other hand, 4f-SAMMY-seq is more consistently classifying all of these Polycomb domains in the “A” compartment (Figure 4a, Supplementary Figure 6c). This was confirmed also in C2C12 (Supplementary Figure 5c) and in human fibroblasts experiments with 50k or 10k cells (Supplementary Figure 7b). Furthermore, in a quantitative analysis, by ordering genomic bins from the most closed (”B”) to the most open (“A”) based on 4f-SAMMY-seq, we observed a linear increase in the percentage of Polycomb state of each region (Figure 4b). Hi-C scatter plot showed a more dispersed distribution and a milder relationship (Figure 4b).

**Figure 4.**
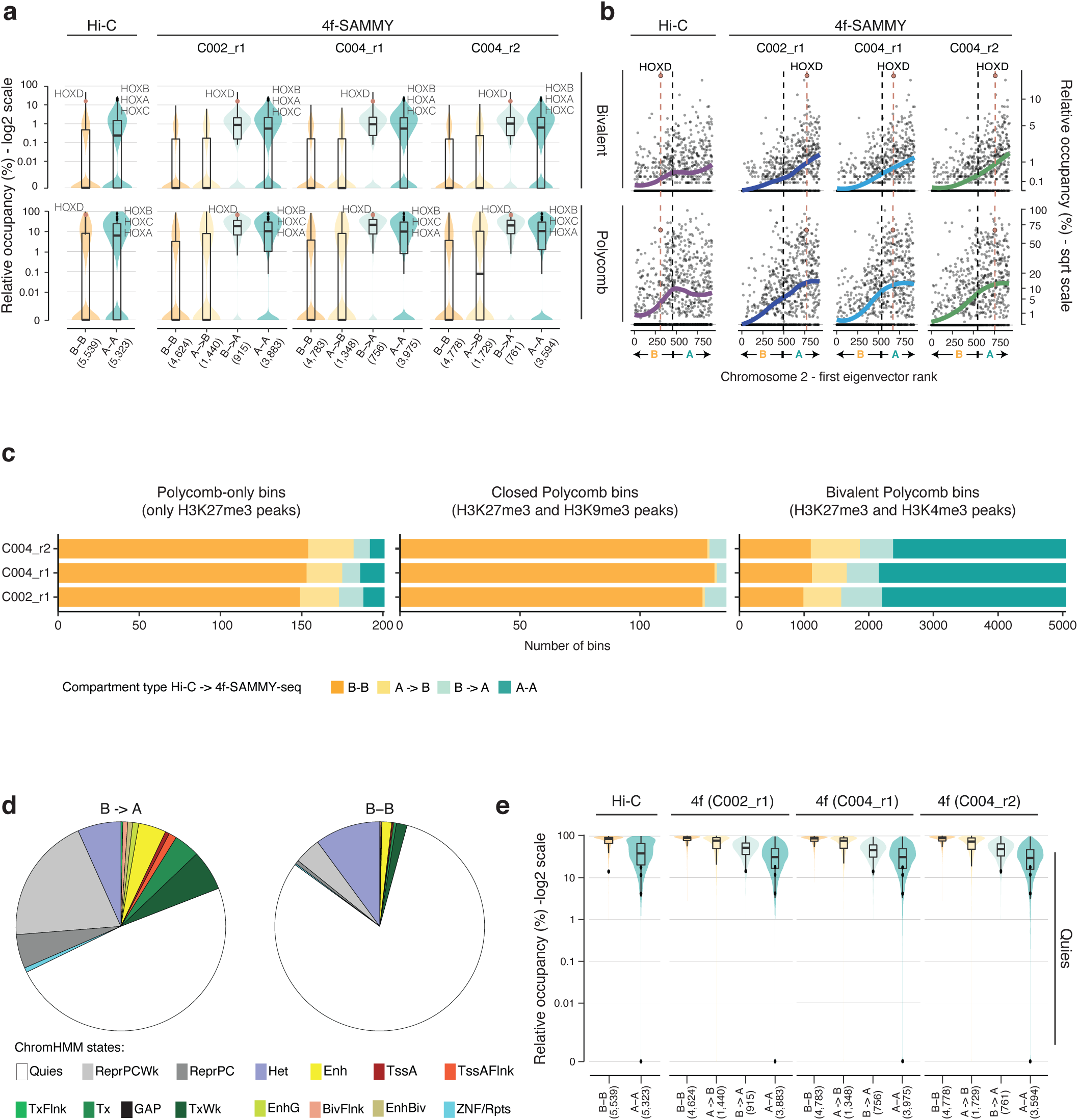
4f-SAMMY-seq based compartments consistently classify Polycomb regulated domains. **a)** Violin and box plots showing for the Hi-C dataset and individual 4f-SAMMY-seq replicates of human fibroblasts (labels on the upper margin) the distribution of Polycomb-regulated chromatin states across genomic bins of the entire genome grouped by compartments classification (”A-A”, “B-B”, “A->B” or “B->A” defined as in Figure 2b). The upper violin and box plots report bivalent Polycomb chromatin states (EnhBiv or BivFlnk) the bottom ones report monovalent repressive Polycomb chromatin states (ReprPC or ReprPCWk). The number of genomic bins in each group is indicated at the bottom in parentheses (x-axis). In the overlaying boxplots, the horizontal line marks the median, the box margins mark the interquartile range (IQR), and whiskers extend up to 1.5 times the IQR. Specific data points associated with HOX gene clusters (black and red dots) are marked to show their positioning across groups and their associated high relative occupancy in Polycomb repressive and bivalent states. **b)** Scatter plots showing the relationship between the rank of the first eigenvector values used to define compartments (ranks on the x-axis) and the relative occupancy by chromatin states plotted as percentage of each bin in squared root (sqrt) scale (y-axis). Each data point shows a 250kb genomic bin from the representative chromosome 2. Black dashed vertical lines mark the transition point between negative and positive values of the first eigenvector, from “B” to “A” compartment. Coloured solid lines highlight the overall trend (lowess regression), for the reference Hi-C dataset (violet line) and for each 4f-SAMMY-seq replicate (blue, light blue and green; labels on upper margins). Dashed salmon vertical lines and circles highlight the HOXD gene cluster bin. Upper plots show the chromatin states occupancy for bivalent Polycomb states and bottom plots monovalent repressive Polycomb states, defined as in panel a. **c)** Barplots reporting the compartments classification for individual replicates for genomic bins overlapping to ChIP-seq peaks for H3K27me3 alone (left barplot), H3K27me3 and H3k9me3 (center barplot), H3K27me3 and H3K4me3 (right barplot). **d)** Pie chart showing the chromatin state composition for discordant (”B->A”) bins overlapping Het chromatin state (left pie chart). The composition of concordant (”B-B”) bins is reported for comparison (right pie chart). **e)** Violin and box plots showing for the Hi-C dataset and for individual 4f-SAMMY-seq replicates (labels on the upper margin) the distribution of the Quiescent (Quies) chromatin states across genomic bins of the entire genome grouped by compartment classification (”A-A”, “B-B”, “A->B” or “B->A” defined as in Figure 2b). The number of genomic bins in each group is indicated at the bottom in parentheses (x-axis). In the overlaying boxplots, the horizontal lines mark the median, the boxes mark the interquartile range (IQR), and whiskers extend up to 1.5 times the IQR.

We further refined this analysis by dividing Polycomb target domains in distinct groups based on ChIP-seq enrichment peaks. Genomic regions enriched for both H3K27me3 and H3K9me3 are generally concordantly classified in B compartment by Hi-C and 4f-SAMMY-seq (Figure 4c). Regions marked exclusively by H3K27me3 are also mostly concordant in B but 4f-SAMMY-seq has an even larger fraction of “A->B”. Whereas when examining bivalent H3K27me3 and H3K4me3 enriched regions we observed a preferential classification in A compartment in both techniques.

As an example of Polycomb-regulated genes, we highlighted the classification of homeobox (HOX) gene clusters (Figure 4a-b). These are a set of developmentally regulated highly conserved gene loci expected to be all enriched in Polycomb-dependent H3K27me3 histone mark in terminally differentiated cells, or paired with H3K4me3 for the bivalent chromatin state (47,48). In humans, 39 HOX genes are organized in 4 distinct chromosomal locations: HOXA, HOXB, HOXC and HOXD gene clusters. While they all show similar chromatin marks, we notice that Hi-C classifies most of them in the “A” compartment, with the sole exception of the HOXD cluster classified in the “B” compartment. Instead, 4f-SAMMY-seq consistently classifies all of them in the “A” compartment (Figure 4a-b and Supplementary Figure 7b).

Among the differences between Hi-C and 4f-SAMMY-seq, the Heterochromatin state emerged for discordant compartment classification (Figure 3b and Supplementary Figure 7a). A further investigation highlighted that actually only a small fraction of this state falls in the A compartment (Supplementary Figure 6c). The discrepancies can be ascribed to the different resolution of the two types of analysis: compartment calls were performed over large genomic bins whereas chromatin state calls defined segments of shorter length. In fact, the discordant bins contain an heterogenous set of active and inactive chromatin states, including a notably larger fraction of Polycomb regulated regions (Figure 4d).

Finally, regions with a “Quies” chromatin state, characterized by weak enrichment in lamina association (Supplementary Figure 6b), are more consistently classified in the “B” compartment based on 4f-SAMMY-seq rather than Hi-C (Figures 3b and 4e).

Overall, these data suggest that 4f-SAMMY-seq derived compartments may provide a consistent reading on the properties of chromatin in terms of their connection to gene expression and chromatin states, including the Polycomb regulated domains, which are more challenging to capture because of their dynamic nature (49).

### 4f-SAMMY-seq also achieves sub-compartments definition

To further dissect the quantitative relation between chromatin marks and compartments, we also computed sub-compartments using an adaptation of the Calder procedure (8) in both Hi-C and 4f-SAMMY-seq (see Methods) (Figure 5a). We called eight sub-compartments spanning from the most active A.1.1 to the most inactive B.2.2 compartment. Both Hi-C and 4f-SAMMY-seq sub-compartments are generally highly concordant in their association with individual chromatin marks (Figure 5b), as confirmed also in the 50k and 10k experiments (Supplementary Figure 7c), as in mouse C2C12 cells (Supplementary Figure 5e). In fact, markers of active chromatin (DNAse-seq, H3K36me3, H3K4me3, H3K27ac and H3K4me1) show the lowest enrichment in the most closed B2.2 sub-compartment and the highest enrichment in the most open A1.1 sub-compartment. Conversely, markers of constitutive heterochromatin (H3K9me3, Lamin AC and Lamin B) show the highest enrichment in B2.2 sub-compartment and a drop of enrichment going towards open sub-compartments.

In line with results observed for A/B compartments classification, also in the sub-compartments analysis the most evident differences involve regions marked by the Polycomb-dependent H3K27me3 histone mark. To this concern, across all datasets we observed: i) higher H3K27me3 enrichment in 4f-SAMMY-seq at A1.1 and A1.2 sub-compartments; ii) enrichment pattern lines for 4f-SAMMY-seq and Hi-C crossing each other at A.2.1 or A.2.2 sub-compartments; iii) lower H3K27me3 enrichment in 4f-SAMMY-seq at B.1.1 and B.1.2 sub-compartments; iv) similar enrichment levels in the most closed sub-compartments (B.2.1 and B.2.2), with slightly lower value in 4f-SAMMY-seq based definition (Figure 5b, Supplementary Figures 5e and 7c). These differences reflect the distinct sensitivity towards the most plastic Polycomb regulated domains as already highlighted in the compartment analysis.

The other main difference on sub-compartments pertains to H3K9me3, for which SAMMY-seq is more specifically confining it to the most inactive B.2.2 sub-compartment, whereas Hi-C has a marked H3K9me3 enrichment also in B.2.1 (Figure 5b, Supplementary Figures 7c). On the other hand, the A.1.1 and A.1.2 sub-compartments from SAMMY-seq are not as depleted of H3K9me3 as those from Hi-C (Figure 5b, Supplementary Figures 7c). In sub-compartment analysis, mouse C2C12 dataset is generally more concordant between Hi-C and 4f-SAMMY-seq, with some differences again for H3K27me3 chromatin mark (Supplementary Figure 5e). Taken together, these results indicate the ability to identify also informative sub-compartments from 4f-SAMMY-seq data, thus achieving a detailed characterization of chromatin compartmentalization in relation to epigenetic marks.

**Figure 5.**
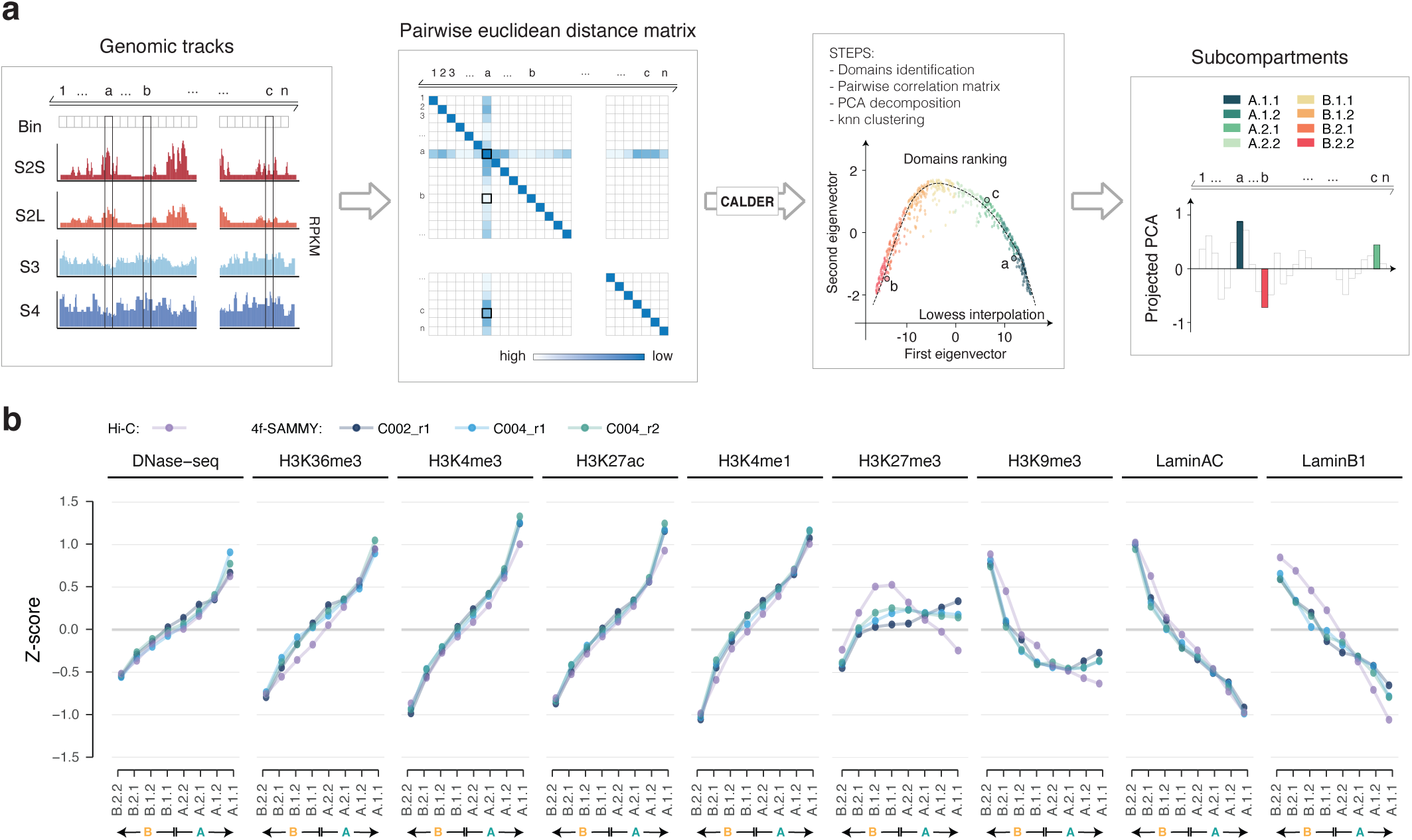
4f-SAMMY-seq allows detailed and reliable reconstruction of sub-compartments. **a)** Schematic illustration of the data analysis workflow to reconstruct chromatin sub-compartments from 4f-SAMMY-seq data. An adaptation of the CALDER procedure is applied on the pairwise Euclidean distance between genomic bins (see Methods). For each pair of genomic bins the Euclidean distance is computed between the vectors of reads coverage across fractions. The method then uses the ranking and clustering of domains (*i.e.*, group of bins) based on the lowess interpolation of the first and second eigenvector of the matrix. From the projected PCA values, the ranking of domains allows discriminating eight sub-compartments from the most compacted one (B.2.2.) to the most accessible one (A.1.1). **b)** Genome-wide mean ChIP-seq IP over INPUT enrichment in human fibroblasts (centred and scaled by chromosome, see Methods) in the eight sub-compartments classification defined by CALDER. Enrichment for DNAse-seq and ChIP-seq datasets is shown for sub-compartments obtained using Hi-C (purple points and lines) and 4f-SAMMY-seq data (blue, light blue and green points and lines for the three replicates). Sub-compartments are sorted from the most compacted (left, B.2.2) to the most accessible one (right, A.1.1).

## DISCUSSION

Chromatin 3D architecture is hierarchically structured on multiple levels to pack and organize the genome inside the nucleus. This emerged as crucial feature for the regulation of transcription, replication, DNA repair and splicing (1). In particular, the physical separation between euchromatin, the more accessible and transcriptionally active part of the genome, and heterochromatin, the more compacted and gene-poor part, is a hallmark of healthy cells. As such, morphological changes in genome organization are emerging as a pathological hallmark, and dysfunctional alterations of chromatin architecture have been described in ageing, cancer and a plethora of other diseases (4,50–52). Heterochromatin can be further divided into facultative heterochromatin, regulated by the Polycomb group of protein (PcG), and constitutive heterochromatin, the latter mainly localized at the nuclear periphery bound to the nuclear lamina (53).

Here we present a novel high-throughput sequencing-based protocol paired with a dedicated bioinformatic data analysis approach to single-handedly identify euchromatin and heterochromatin domains, as well as mapping their 3D compartmentalization inside the cell nucleus. The new 4f-SAMMY-seq method builds on our previously published 3f-SAMMY-seq technique, for which we have already demonstrated the ability to detect lamina-associated heterochromatin domains dynamics (16,19). Here we developed a protocol with two crucial novel steps including a milder partial endonuclease digestion of accessible chromatin and size separation of the resulting DNA fragments, to achieve higher resolution in mapping accessible chromatin regions (Figure 1, Supplementary Figure 2). We explored multiple conditions, finely modulating the DNAse and genomic DNA amount to obtain, with a unique protocol, the higher correlations with specific marks of open and closed chromatin (Supplementary Figure 2a and 4a-c).

We then reasoned on the differences between our chromatin fractionation-based method and Hi-C, that uses frequency of contacts between genomic loci to infer chromatin compartmentalization. On the other hand, SAMMY-seq derived methods rely on the biochemical isolation of distinct chromatin fractions, then assessing the differential enrichment of genomic sequences across them. Genomic loci close to each other generally have the same epigenetic chromatin state, thus they are exposed to a similar mixture of DNA, RNA, proteins and epigenetic modifications constituting the local biochemical environment. We hypothesized that we could reconstruct chromatin compartmentalization by directly measuring the similarity of genomic loci distribution across 4f-SAMMY-seq fractions, which capture the biochemical properties of chromatin in terms of accessibility and solubility.

Thus, we developed and present here a bioinformatic approach to reconstruct the large-scale chromatin 3D organization in compartments and sub-compartments (Figure 2 and 5). The results are similar to the ones derived from Hi-C, but with some notable differences.

Our analyses indicate that 4f-SAMMY-seq consistently captures specific dynamic chromatin domains such as Polycomb regulated states (Figures 3 and 4). Polycomb targets are heterogeneous from a functional and regulatory dynamics point of view (49). Different combinations of Polycomb complexes can modulate the transcriptional output as well as the context specific regulation of individual target loci (54). They have a central role in many physiological processes and they are altered in several human diseases (55,56). Their role in determining chromatin 3D architecture was already established years ago first in Drosophila (57) then in mammalian cells (58,59). Polycomb deposited histone marks (H3K27me3) are often found at the boundary of LADs (42,60,61), and interfering with either LADs or Polycomb regulation has been shown to have an effect on each other (13,62). On the other hand, Polycomb targets include bivalent loci, genes simultaneously marked by active and inactive histone modifications priming them for a rapid transcriptional activation (47). These features confirm that Polycomb targets are a very heterogeneous and dynamic portion of chromatin, in addition to be crucial for genome 3D organization. For these reason Polycomb regulated regions are especially challenging for the all methods used to study chromatin structure.

Hi-C historically classified Polycomb-regulated domains as belonging to the “A” compartment (5), or in an “intermediate” compartment, yet mostly associated to positive eigenvector values (4). In this regard, we found that SAMMY-seq-based compartments provide a more consistent and concordant classification of Polycomb targets, including the highly dynamic bivalent genes (Figure 4a-b).

A separate discussion is required for sub-compartments, that are historically identified with a different bioinformatic approach. Namely, sub-compartments were first identified based only on trans (i.e. inter-chromosomal) interactions (7), then their association to H3K27me3 was observed specifically for an intermediate B1 compartment, but with some degree of interaction also with the active A1 compartment as reported in the original article. Later approaches also adopted only trans-interactions for the sub-compartments definition (63), with the notable exception of CALDER (8), that is using both cis and trans interactions, unbiasedly placing H3K27me3 enriched domains in sub-compartments of the “B” group. We used this algorithm in our comparison between SAMMY-derived and Hi-C-derived sub-compartments and we found again some specific, yet consistent, differences for H3K27me3 enrichment distribution (Figure 5). In SAMMY-seq analysis we observed a higher H3K27me3 enrichment in A-type sub-compartments (Figure 5, Supplementary 7). Interestingly, in C2C12 dataset, although we confirm PcG domains assignment in A compartment (Supplementary Figure 5b-c), the trend of sub-compartment analysis is closer to Hi-C.

It is worth remarking that Hi-C and SAMMY-seq are based on different principles, thus capturing the nuances of the dynamic Polycomb epigenetically regulated regions with different quantitative patterns across compartments or sub-compartments, that could be dependent on the cell type. This should be taken into consideration for future applications of the techniques. In this context, the 4fSAMMY-seq is a versatile technology with unique characteristics that enable its applicability to several experimental models (Supplementary Table 2).

First of all, with 4f-SAMMY-seq protocol it is possible to simultaneously analyse chromatin accessibility and compartments, which normally require distinct genomic techniques. Moreover, while the standard Hi-C protocol requires at least 1 million cells, here we showed that our method works on as little as 10,000 cells, thus opening unprecedented opportunities in terms of applications to small biological samples (e.g. patients biopsies or rare cell populations). Moreover, to achieve high resolution, Hi-C libraries are often sequenced to more than one billion reads depth, whereas with 4f-SAMMY-seq we can use as little as 25 million reads per chromatin fraction (see Methods). Finally, the 4f-SAMMY-seq experimental protocol requires less than three hours in sample manipulation time, as opposed to at least two days for Hi-C. For all of these practical advantages, applying 4f-SAMMY-seq will enable unparalleled scalability and applicability in the study of chromatin compartment dynamics.

## DATA AVAILABILITY

The high-throughput sequencing data generated for this study are available in the NCBI GEO database with accession number “GSE226718”.

The bioinformatic pipeline along with a step-by-step tutorial explaining how to process SAMMY-seq data is available in the Github repository https://github.com/Clockris/SAMMY-method_4f.

## AUTHOR CONTRIBUTIONS

FL, VR, FGo, PS, RQ and CL performed the experiments. CP and ESa performed computational data analyses. KP contributed to the initial conceptualization of compartment analysis with 4f-SAMMY-seq. KP, FL, GL and GG contributed to specific data analyses. EDPS, IT, EP and ESe contributed to preliminary data analyses and testing of analyses procedures. LR, FGe and VV contributed to high-throughput sequencing. FL, CP, ESa, CL and FF wrote the manuscript. CL and FF jointly supervised the work and designed the study. All authors reviewed and approved the manuscript for submission.

## ACKNOWLEDGEMENTS

We thank Beatrice Bodega, Gioacchino Natoli, Daniel Jost, Marina Lusic, Maria Vivo, Yinxiu Zhan, Vittore Scolari, Federica Marasca and all members of our laboratories for critical feedback on earlier versions of the manuscript. We acknowledge the use of dbGaP data from the study with accession number phs000791.v2.p1 (part of the Roadmap Epigenomics program). We thank Margherita Mutarelli (CNR) for NextFlow porting of SAMMY-seq analysis pipeline. We acknowledge support from the Genomics Unit (Thelma Capra) of the European Institute of Oncology (IEO), Department of Experimental Oncology, from the Sequencing Unit (Giulia Passignani) of Fondazione IRCCS Ca’ Granda-Ospedale Maggiore Policlinico and Istituto Nazionale Genetica Molecolare (INGM) (Marco Ghilotti).

---

**Supplementary Figure 1.**
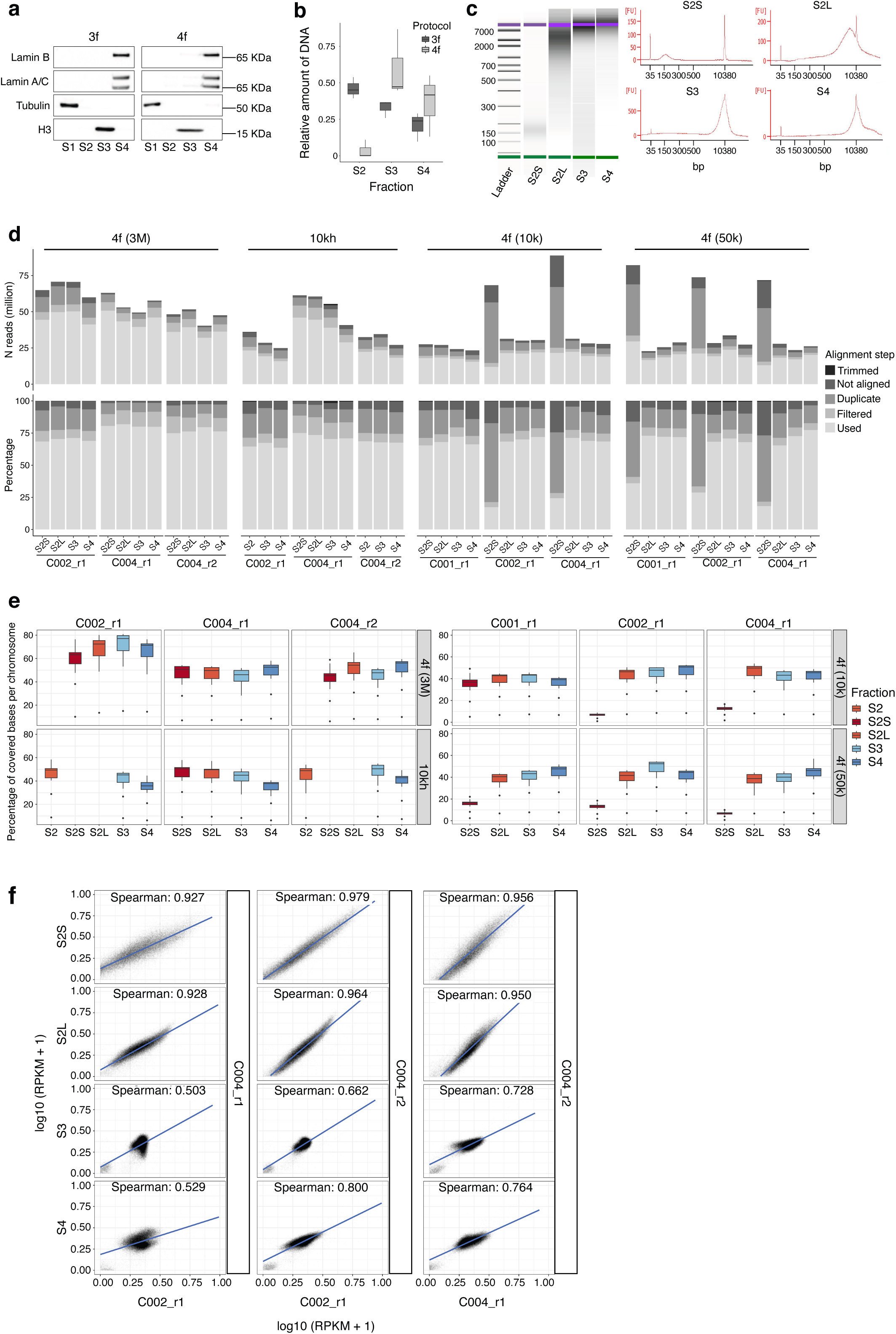
Quality controls on SAMMY-seq experimental procedure. **a)** Representative western blots of chromatin fractions obtained with “3f” and “4f” SAMMY-seq protocols. In both cases we find soluble tubulin in S1 fraction, histone H3 in S3 fraction and lamins in S4 fraction. **b)** Relative abundance (y-axis) of DNA extracted from S2, S3 and S4 fractions, computed as ratio over their sum, in “3f” and “4f” SAMMY-seq protocols (dark and light grey, respectively). In the boxplots, the horizontal lines mark the median, the boxes mark the interquartile range (IQR) and whiskers extend up to 1.5 times the IQR. Relative abundance of DNA was evaluated over 3 independent biological replicates for each protocol. **c)** Representative bioanalyzer electropherograms of individual 4f-SAMMY-seq fractions showing the size distribution of DNA fragments before sonication. **d)** Stacked barplots for the number of sequencing reads obtained for each sample and chromatin fraction, divided in trimmed, not aligned, duplicates, filtered (discarded) and used reads retained for downstream analyses. See also the associated Supplementary Table 1. Individual replicates are reported for 4f-SAMMY-seq experiments on human fibroblasts (3M, 3 million, indicating the starting number of cells), as well as for the 10kh-SAMMY-seq protocol variant, and the scale-down experiments with 10k or 50k starting number of cells. **e)** Boxplots summarizing coverage across chromosomes for each sample and fraction. The percentage of each chromosome covered by at least one read is reported on the y-axis. In the boxplots, the horizontal lines mark the median, the boxes mark the interquartile range (IQR) and whiskers extend up to 1.5 times the IQR. Individual replicates are reported for 4f-SAMMY-seq experiments, as well as for the 10kh-SAMMY-seq protocol variant, and the scale-down experiments with 10k or 50k starting number of cells. **f)** Scatter plots of reads distribution profiles (RPKM normalized coverage over 50kb bins) for individual fractions of 4f-SAMMY-seq replicates. Fractions labels indicated on individual rows (labels on the left) and the replicates compared in each plot are indicated on bottom and right-side axes.

**Supplementary Figure 2.**
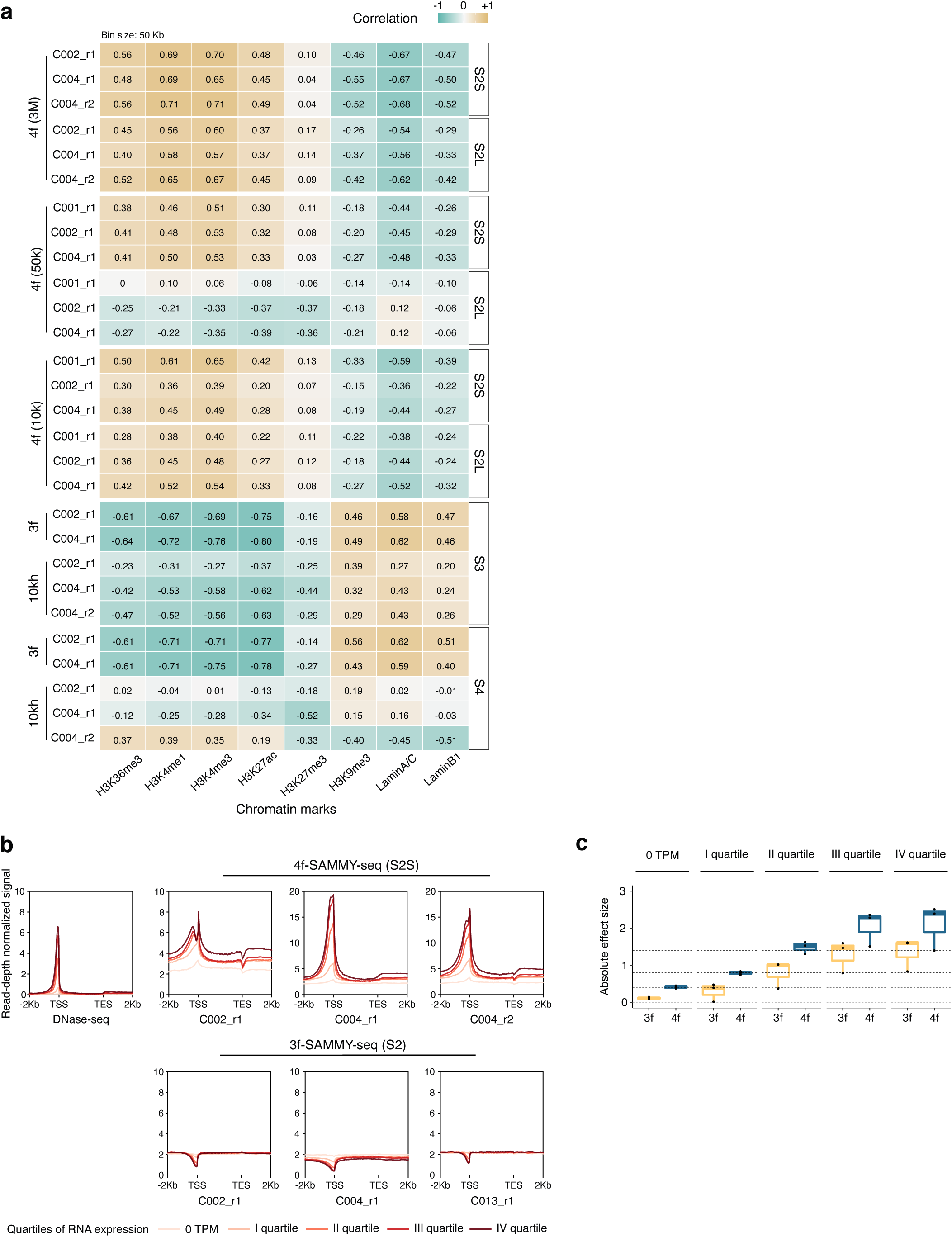
4f-SAMMY-seq and 10kh-SAMMY-seq recapitulate open and closed chromatin domains. **a)** Genome-wide Spearman correlation between reads distribution profiles for individual selected SAMMY-seq chromatin fractions and ChIP-seq enrichment profiles for histone marks and lamin proteins (x-axis labels) from human fibroblasts. The labels on the left side indicate the protocol versions for 3f-SAMMY-seq (3f), 4f-SAMMY-seq (4f) starting from 3million cells (3M), 4f-SAMMY-seq starting from 50,000 or 10,000 cells (50k and 10k, respectively), 10kh-SAMMY-seq (10kh). The labels for individual chromatin fractions (S2S, S2L, S3 and S4) are reported on the right. A row for each replicate is shown and correlation values are reported as numbers and as colour gradient. **b)** Gene centred meta-profiles for reads distribution profiles of a reference DNase-seq sample (foreskin human fibroblasts from ENCODE, aliases: roadmap-epigenomics:E055) and individual replicates of 4f-SAMMY-seq S2S fraction and 3f-SAMMY-seq S2 fraction. Genes are divided by quartiles of expression level (RPKM) and a meta-profile is drawn for each quartile, as well as for genes with zero RPKM (no reads), as measured by RNA-seq. On the x-axis, the coordinates for the relative genomic position around the rescaled gene body are reported. **c)** Paired effect size was computed with Cohen’s *d* method for the difference in reads coverage at TSS and at an upstream region (-2Kb) used as reference background. The absolute effect size value is reported in the boxplot for each replicate (individual dots reported in the plot) and for each expression quartile (labels on top). Data are shown separately for the 3f-SAMMY-seq and 4f-SAMMY-seq experiments (yellow and blue boxes, respectively) from the corresponding panel b. Dashed lines demarcate standard Cohen’s *d* cut-offs indicating none, small (>0.2), medium (>0.4), large (>0.8) and very large (>1.4) effect size.

**Supplementary Figure 3.**
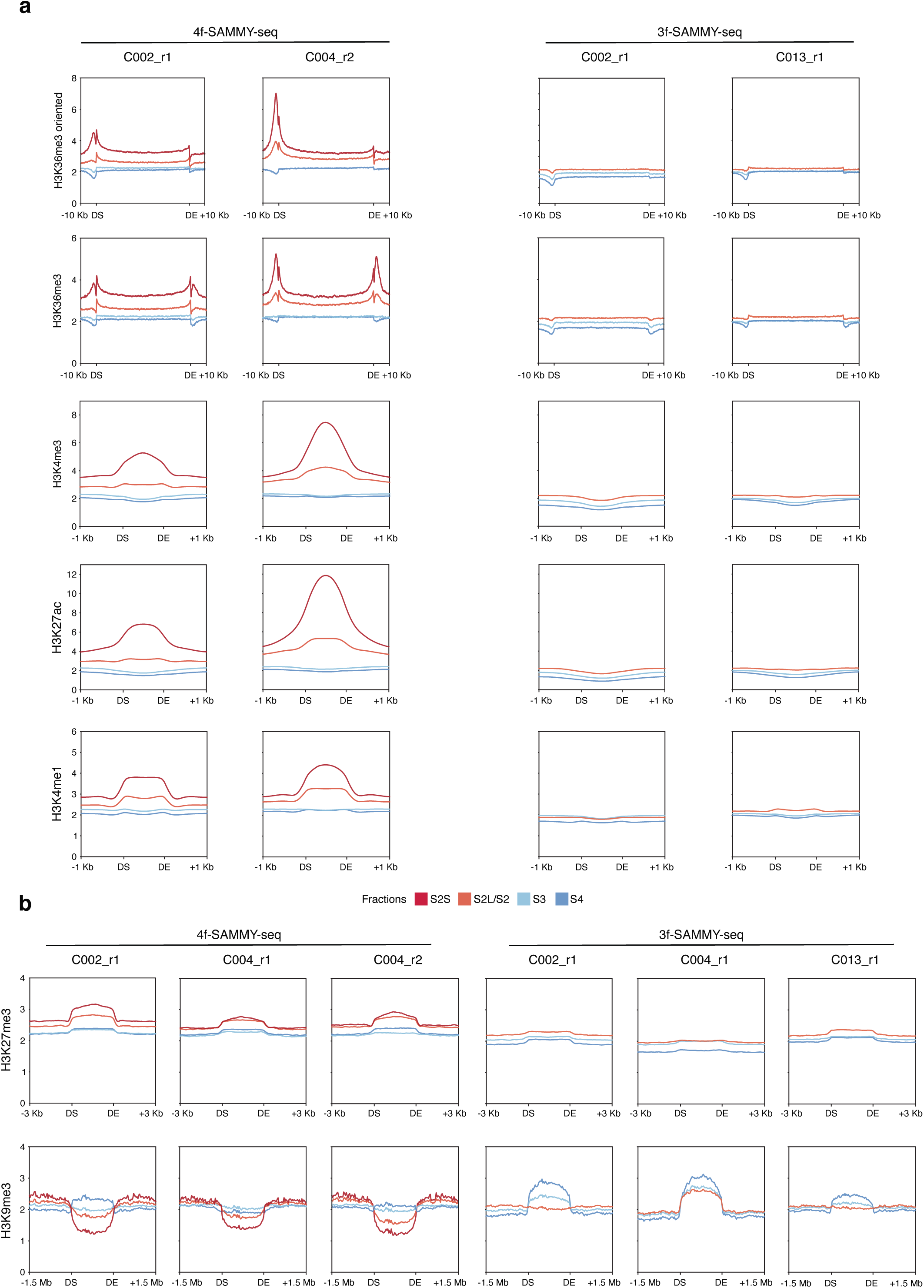
Chromatin fractions read profiles recapitulate epigenetic marks. **a)** Reads distribution meta-profiles for individual “4f” or “3f” SAMMY-seq fractions (labels on the top border). The average reads distribution profiles are computed over chromatin domains marked by enrichment peaks of specific histone marks (indicated on the left side of each row). The domain start (DS) and domain end (DE) are indicated on the x-axis along with flanking regions coordinates. For H3K36me3 mark, we reported also the meta-profile obtained by orienting the domains according to the corresponding gene’s transcribed strand. **b)** Reads distribution meta-profiles following the same colouring and labelling conventions adopted in panel a, but referring to domains enriched for histone marks associated to inactive chromatin: H3K27me3 and H3K9me3.

**Supplementary Figure 4.**
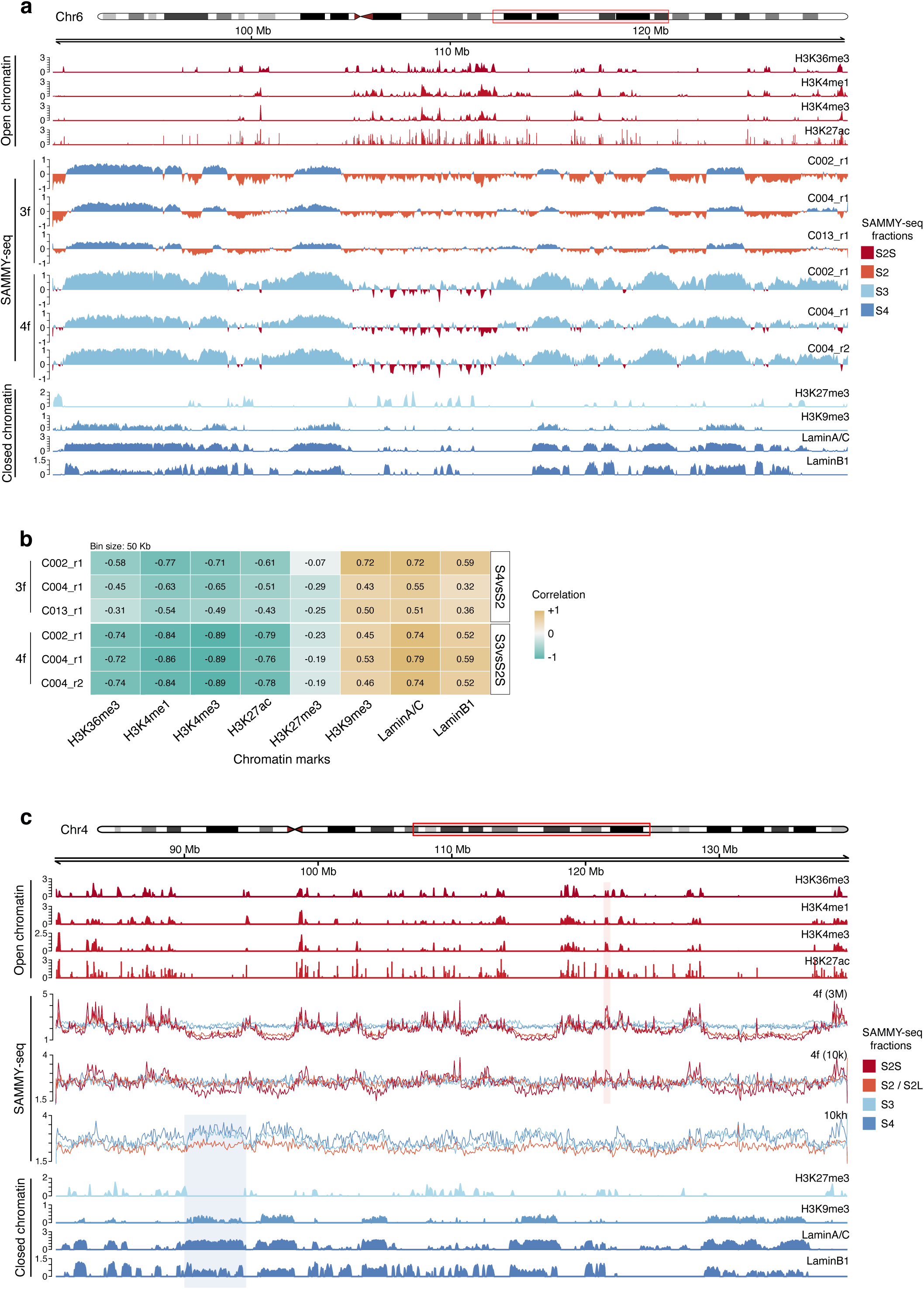
Relative comparison of chromatin fractions recapitulate heterochromatin regions. **a)** Representative genomic region of (chr6:62,000,000-130,000,000) showing tracks for SAMMY-seq and chromatin marks in human fibroblasts. From top to bottom: open chromatin marks ChIP-seq enrichment profiles for H3K36me3, H3K4me1, H3K4me3, H3K27ac; relative enrichment of closed (S3 or S4) over accessible (S2 or S2S) (see color legend) chromatin fractions from “3f” and “4f” SAMMY-seq in all replicates; closed chromatin marks ChIP-seq enrichment profiles for H3K27me3, H3K9me3, Lamin A/C, Lamin B1. **b)** Genome-wide Spearman correlation between relative enrichment profiles of closed (S3 or S4) over accessible (S2 or S2S) (labels on the right) chromatin fractions from “3f” and “4f” SAMMY-seq protocol variants (labels on the left) for individual replicates. **c)** Representative genomic region of (chr4:80,000,000-140,000,000) showing tracks for SAMMY-seq and chromatin marks in human fibroblasts. From top to bottom: open chromatin marks ChIP-seq enrichment profiles for H3K36me3, H3K4me1, H3K4me3, H3K27ac; reads distribution profiles for individual fractions of a representative replicate of 4f-SAMMY-seq on 3M (C004_r2) and 10k cells (C001_r1), as well as 10kh-SAMMY-seq (C002_r1); closed chromatin marks ChIP-seq enrichment profiles for H3K27me3, H3K9me3, Lamin A/C, Lamin B1. The shaded areas mark two examples of regions showing enrichment for closed (blue) or open (red) chromatin marks, with corresponding enrichment patterns visible in ChIP-seq as well as in SAMMY-seq fractions.

**Supplementary Figure 5.**
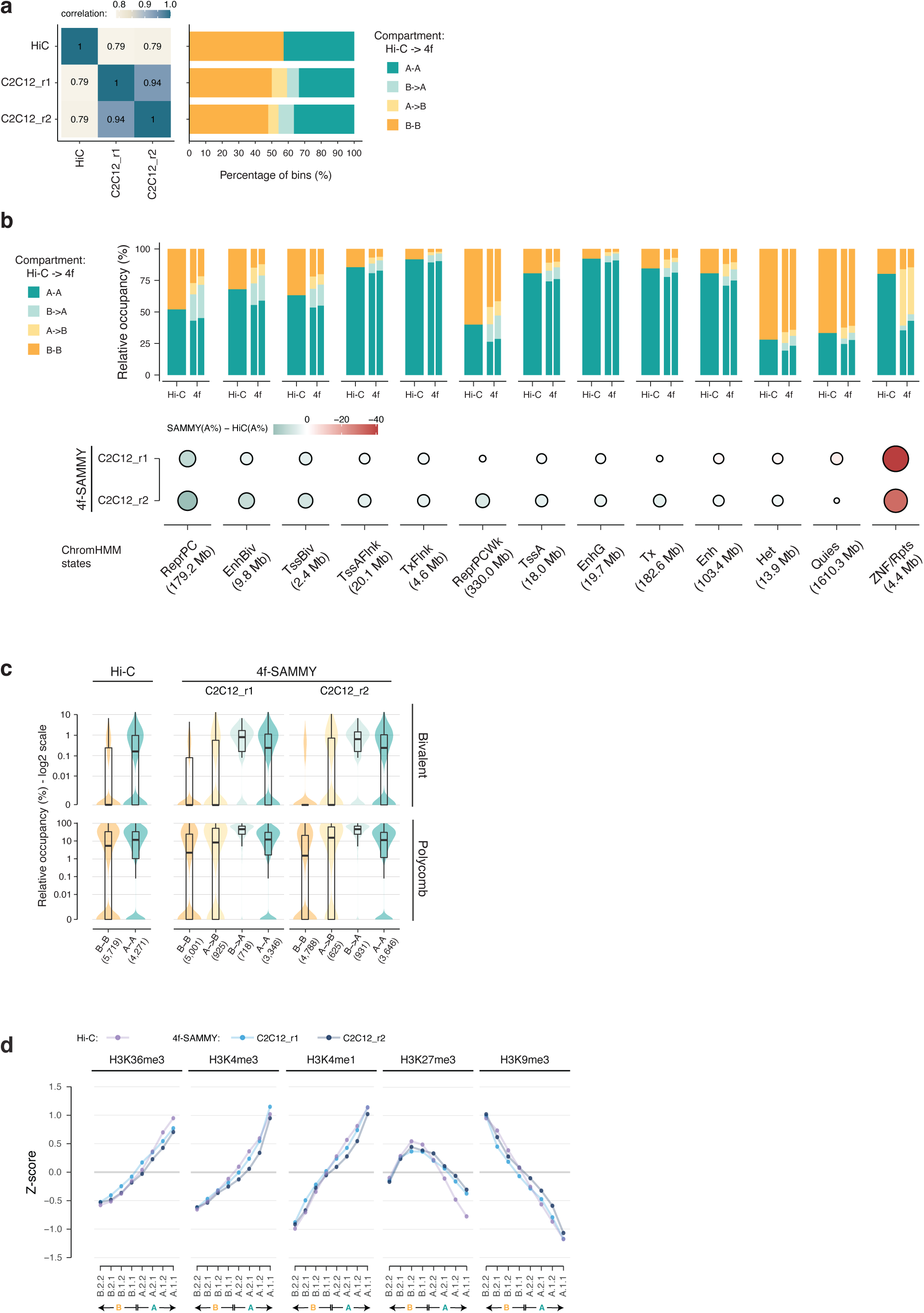
4f-SAMMY-seq based compartments and sub-compartments on mouse C2C12 cells. **a)** Genome-wide pairwise Pearson correlation of chromatin compartments eigenvectors (250kb size bins) defined by Hi-C and 4f-SAMMY-seq in mouse C2C12 cells with individual replicates is reported on the left. For each sample, the stacked barplot on the right shows the relative distribution (percentage) of genomic bins associated with concordant (”A-A” or “B-B”) and discordant (”A->B” or “B->A”) compartment classification with respect to Hi-C. The classification is reported using the same colouring and naming convention adopted in Figure 2b. **b)** Distribution of “A” and “B” compartments (from Hi-C and 4f-SAMMY-seq) across multiple chromatin states from mouse C2C12 myoblasts. The stacked barplots show the percentage of regions associated to concordant (”A-A” or “B-B”) and discordant (”A->B” or “B->A”) compartments classification for each chromatin state (labels on the bottom margin indicating chromatin states and their total size). The barplot for 4f-SAMMY-seq compartments distribution is divided in two parts to report the specific results for the two replicates (C2C12_r1 and C2C12_r2, respectively). Below the barplot, the dot plot reports for each chromatin state the relative occupancy of 4f-SAMMY-seq vs Hi-C based compartments, computed as the difference in “A” compartment percentage. For each chromatin state (columns, labels and total size for each state at the bottom) positive values (green gradient) indicate a higher percentage of “A” compartment in 4f-SAMMY-seq based classification, whereas negative values (red gradient) indicate a higher percentage of “A” compartment based on Hi-C classification (i.e. higher “B” percentage based on 4f-SAMMY-seq). The size of each dot is proportional to the absolute value in the percentage difference. Chromatin states are ordered from left to right based on the average percentage difference across the 4f-SAMMY-seq replicates (on the rows). **c)** Violin and box plots showing for the mouse C2C12 Hi-C dataset and individual 4f-SAMMY-seq replicates (labels on the upper margin) the distribution of Polycomb regulated chromatin states across genomic bins of the entire genome grouped by compartments classification (”A-A”, “B-B”, “A->B” or “B->A” defined as above). The upper violin plot is for bivalent Polycomb states (TssBiv or EnhBiv) the bottom one for monovalent repressive Polycomb states (ReprPC or ReprPCWk). The number of genomic bins in each group is indicated at the bottom in parentheses (x-axis). In the overlaying boxplots, the horizontal lines mark the median, the boxes margins mark the interquartile range (IQR), and whiskers extend up to 1.5 times the IQR. Specific data points associated with HOX gene clusters are marked to show their positioning across groups and their associated high relative occupancy in Polycomb repressive and bivalent states. **d)** Genome-wide mean ChIP-seq IP over INPUT enrichment in C2C12 cells (centred and scaled by chromosome, see Methods) in the eight sub-compartments classification defined by CALDER. Enrichment for ChIP-seq datasets is shown for sub-compartments obtained using Hi-C (purple points and lines) and 4f-SAMMY-seq data (blue and light blue points and lines for the two replicates). Sub-compartments are sorted from the most compacted (left, B.2.2) to the most accessible one (right, A.1.1).

**Supplementary Figure 6.**
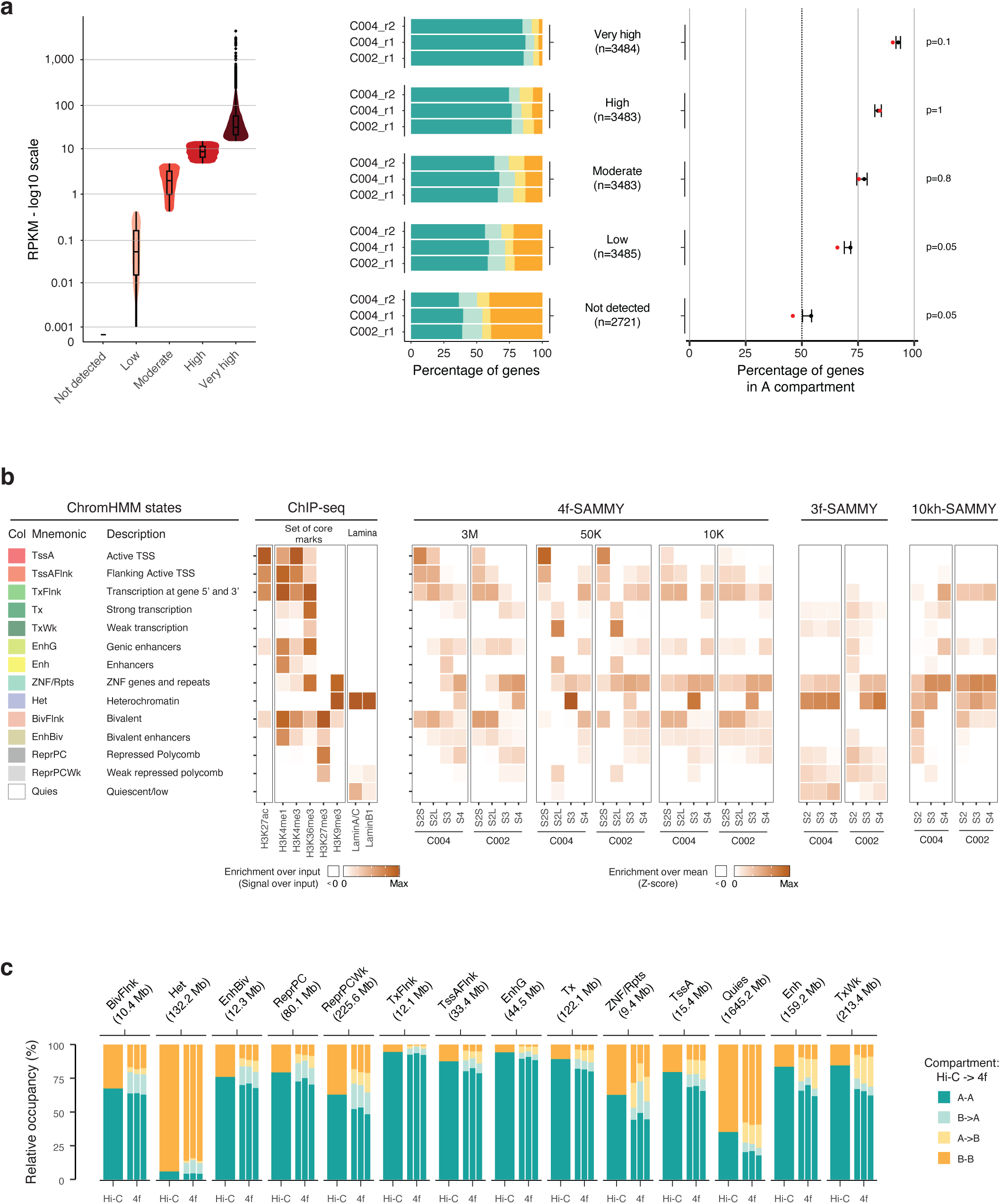
4f-SAMMY-seq chromatin fractions and chromatin marks association to chromatin states. **a)** Compartments classification comparison with gene expression status. Genes are divided by quartiles of expression level (RPKM values from Roadmap Epigenomic E055 sample) and labelled as “Low”, “Moderate”, “High” and “Very high” according to their expression level (violin and box plot on the left). Genes with zero RPKM (no reads as measured by RNA-seq) are assigned to the “Not detected” group. The stacked barplot shows for each 4f-SAMMY-seq replicate the relative distribution of genes across concordant and discordant compartments classifications with respect to Hi-C compartments (50kb size bins - genes assigned to bins based on the TSS position). The chromatin compartment classification is reported using the same colouring and naming convention adopted in Figure 2b. In the dot and whiskers plot on the right we show the percentage of genes mapped in A compartment by Hi-C (red dot) and by each 4f-SAMMY-seq replicates (whiskers for maximum and minimum range, dot for the median replicate). The significance of differences was tested with a one-sided t-test. **b)** Epigenetic marks enrichment signatures associated to chromHMM chromatin states. We called chromatin states on a compendium of proprietary and public ChIP-seq datasets and assigned labels to maximize comparability to the chromatin states annotations adopted by the Roadmap Epigenomic consortium. Starting from the left: the colour code, the name and the description of the chromatin states, the heatmaps representing the enrichment over the input for H3K27ac, H3K4me1, H3K4me3, H3K36me3, H3K27me3 and H3K9me3, lamin A/C and B1, the heatmaps representing the enrichment over the mean (z-score) of individual SAMMY-seq fractions for protocol versions “4f” (3M, 50K and 10K), “3f” and “10kh” with two representative replicates for each of them. The histone marks originally used in the chromHMM definition of states are grouped in the “core marks” quadrant in the figure. **c)** The stacked barplots show the percentage of regions associated with concordant (”A-A” or “B-B”) and discordant (”A->B” or “B->A”) compartment classification for each chromatin state (labels on the upper margin indicating chromatin states and their total size). The chromatin compartments classification is reported using the same colouring convention adopted in Figure 2b. The barplot for 4f-SAMMY-seq compartments distribution is divided in three parts to report the specific result for the three replicates (C002_r1, C004_r1 and C004_r2, respectively). The chromatin states are ordered from left to right based on the average difference between Hi-C and SAMMY-seq based compartments.

**Supplementary Figure 7.**
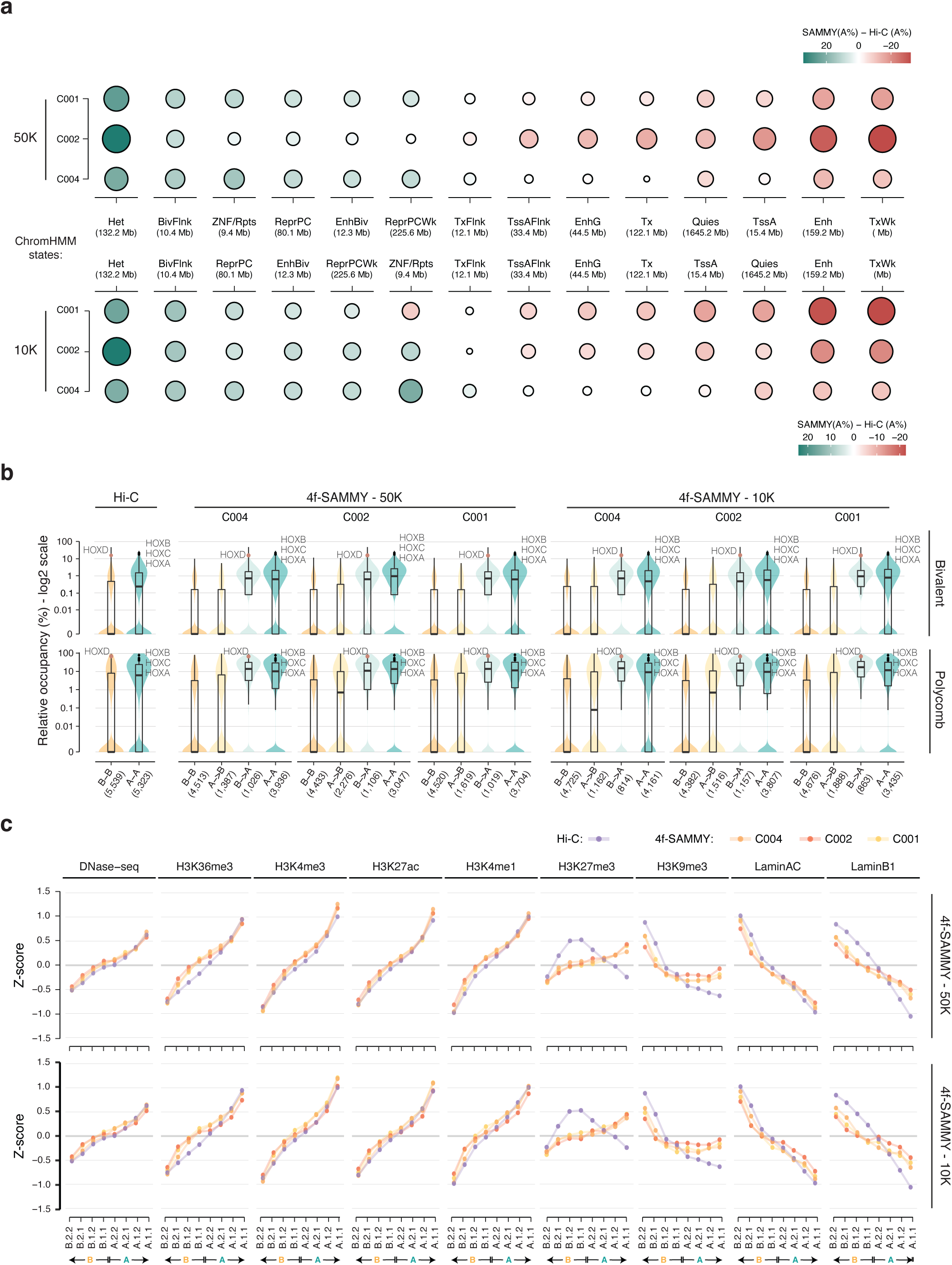
Compartments and sub-compartments analysis on low input 4f-SAMMY-seq. **a)** Relative occupancy of 4f-SAMMY-seq (starting from either 50k or 10k cells as indicated on the left side labels) vs Hi-C based compartments computed as the difference in “A” compartment percentage. For each chromatin state (columns, labels and total size for each state in the central row) positive values (green gradient) indicate a higher percentage of “A” compartment in 4f-SAMMY-seq based classification, whereas negative values (red gradient) indicate a higher percentage of “A” compartment based on Hi-C classification (*i.e.* higher “B” percentage based on 4f-SAMMY-seq). The size of each dot is proportional to the absolute value in the percentage difference. Chromatin states are ordered from left to right based on the average percentage difference across the three 4f-SAMMY-seq replicates (on the three rows). **b)** Violin and box plots showing for the Hi-C dataset and for individual 4f-SAMMY-seq replicates starting from either 50k or 10k cells (labels on the upper margin) the distribution of Polycomb regulated chromatin states across genomic bins of the entire genome grouped by compartments classification (”A-A”, “B-B”, “A->B” or “B->A” defined as in Figure 2b). The upper violin plot is for bivalent Polycomb states (EnhBiv or BivFlnk) the bottom one for monovalent repressive Polycomb chromatin states (ReprPC or ReprPCWk). The number of genomic bins in each group is indicated at the bottom in parentheses (x-axis). In the overlaying boxplots, the horizontal lines mark the median, the boxes margins mark the interquartile range (IQR), and whiskers extend up to 1.5 times the IQR. Specific data points associated with HOX gene clusters are marked to show their positioning across groups and their associated high relative occupancy in Polycomb repressive and bivalent states. **c)** Genome-wide mean ChIP-seq IP over INPUT enrichment in human fibroblasts (centred and scaled by chromosome, see Methods) in the eight sub-compartments classification defined by CALDER. Enrichment for DNAse-seq and ChIP-seq datasets is shown for sub-compartments obtained using Hi-C (purple points and lines) and 4f-SAMMY-seq data obtained starting from 50K and 10K (red, orange and yellow points and lines for the three replicates). Sub-compartments are sorted from the most compacted (left, B.2.2) to the most accessible one (right, A.1.1).

